# *Plasmodium* condensin core subunits (SMC2/SMC4) mediate atypical mitosis and are essential for parasite proliferation and transmission

**DOI:** 10.1101/674028

**Authors:** Rajan Pandey, Steven Abel, Matthew Boucher, Richard J. Wall, Mohammad Zeeshan, Edward Rea, Aline Freville, Xueqing Maggie Lu, Declan Brady, Emilie Daniel, Rebecca R. Stanway, Sally Wheatley, Gayani Batugedara, Thomas Hollin, Andrew R. Bottrill, Dinesh Gupta, Anthony A. Holder, Karine G. Le Roch, Rita Tewari

**Affiliations:** School of Life Sciences, Queens Medical Centre, University of Nottingham, Nottingham, NG7 2UH, UK; Department of Molecular, Cell and Systems Biology, University of California Riverside, 900,University Ave, Riverside, CA, 92521, USA; Wellcome Trust Centre for Anti-Infectives Research, School of Life Sciences, University of Dundee, Dundee, DD1 5EH, UK; Institute of Cell Biology, University of Bern, Bern 3012, Switzerland; School of Life Sciences, Gibbet Hill Campus, University of Warwick, Coventry, CV4 7AL, UK; Translational Bioinformatics Group, International Center for Genetic Engineering and Biotechnology, New Delhi, 110067, India; Malaria Parasitology Laboratory, The Francis Crick Institute, London, NW1 1AT, UK

## Abstract

Condensin is a multi-subunit protein complex regulating chromosome condensation and segregation during cell division. In *Plasmodium* spp., the causative agent of malaria, cell division is atypical and the role of condensin is unclear. Here we examine the role of SMC2 and SMC4, the core subunits of condensin, during endomitosis in schizogony and endoreduplication in male gametogenesis. During early schizogony SMC2/SMC4 localize to a distinct focus, identified as the centromeres by NDC80 fluorescence and ChIP-seq analyses, but do not form condensin I or II complexes. In mature schizonts and during male gametogenesis, there is a diffuse SMC2/SMC4 distribution on chromosomes and in the nucleus, and both condensin I and II complexes form at these stages. Knockdown of *smc2* and *smc4* gene expression revealed essential roles in parasite proliferation and transmission. The condensin core subunits (SMC2/SMC4) form different complexes and may have distinct functions at various stages of the parasite life cycle.

## Introduction

Cellular proliferation in eukaryotes requires chromosome replication and segregation, followed by cell division, to ensure that daughter cells have identical copies of the genome. During classical open mitosis in many eukaryotes, chromosome condensation, centrosome migration and formation of the mitotic spindle are followed by dissolution of the nuclear envelope (Guttinger et al., 2009). In contrast, in some unicellular organisms such as the budding yeast, *Saccharomyces cerevisiae*, mitosis is closed: the nuclear membrane remains intact and chromosomes are separated by spindles assembled within the nucleus (Sazer et al., 2014). The mechanisms and the various regulatory molecules involved in cell division have been well studied in many eukaryotes. The cell division regulators include cyclins, cyclin-dependent kinases (CDKs), components of the anaphase promoting complex (APC), and other protein kinases and phosphatases (Chang et al., 2014; Fisher et al., 2012; Harashima et al., 2013).

An essential component of chromosome dynamics is a family of ‘Structural Maintenance of Chromosomes’ proteins, originally described in budding yeast as ‘stability of minichromosomes’ (SMC) proteins, which are implicated in chromosome segregation and condensation (Hirano, 2016; Uhlmann, 2016). Most eukaryotes have at least six genes encoding SMC proteins (each 110-170 kDa, with a central hinge region and N and C-terminal globular domains with Walker A and Walker B motifs forming the ATPase head domain). The six SMCs can be classified as subunits of condensin (SMC2 and SMC4, required for chromosomal condensation), cohesin (SMC1 and SMC3, required for chromosomal segregation) and the SMC5-SMC6 complex (involved in DNA repair and homologous recombination) (Hirano, 2016; Uhlmann, 2016).

Higher eukaryotic organisms have two different condensin complexes: condensin I and condensin II, whereas many single-celled organisms such as yeast have only one condensin complex. SMC2 and SMC4 form the core structure for both condensin I and condensin II in higher eukaryotes (Hirano, 2016), and interact with three additional non-SMC components, including one kleisin (Schleiffer et al., 2003) and two heat protein subunits (Neuwald and Hirano, 2000). Kleisin Iγ (CAP-H), Heat IA (CAP-D2) and Heat IB (CAP-G) form the condensin I complex, whereas Kleisin IIβ (CAP-H2), Heat IIA (CAP-D3) and Heat IIB (CAP-G2) form the condensin II complex (Hirano, 2016; Uhlmann, 2016) (**Figure 1A**). Electron microscopy and protein-protein interaction studies have revealed the characteristic architecture and geometry of condensin complexes (Anderson et al., 2002; Onn et al., 2007). Condensin plays a vital role in cell division processes such as chromosomal condensation, correct folding and organization of chromosomes prior to anaphase, and proper chromosome segregation and separation (Hirano, 2016; Kschonsak et al., 2017; Ono et al., 2013; Rawlings et al., 2011; Uhlmann, 2016). Both SMC and non-SMC components are necessary for full function; for example, chromosomal condensation is not observed in the absence of kleisin, showing its critical role for complex formation and condensation (Cuylen et al., 2011; Rawlings et al., 2011).

**Figure 1:**
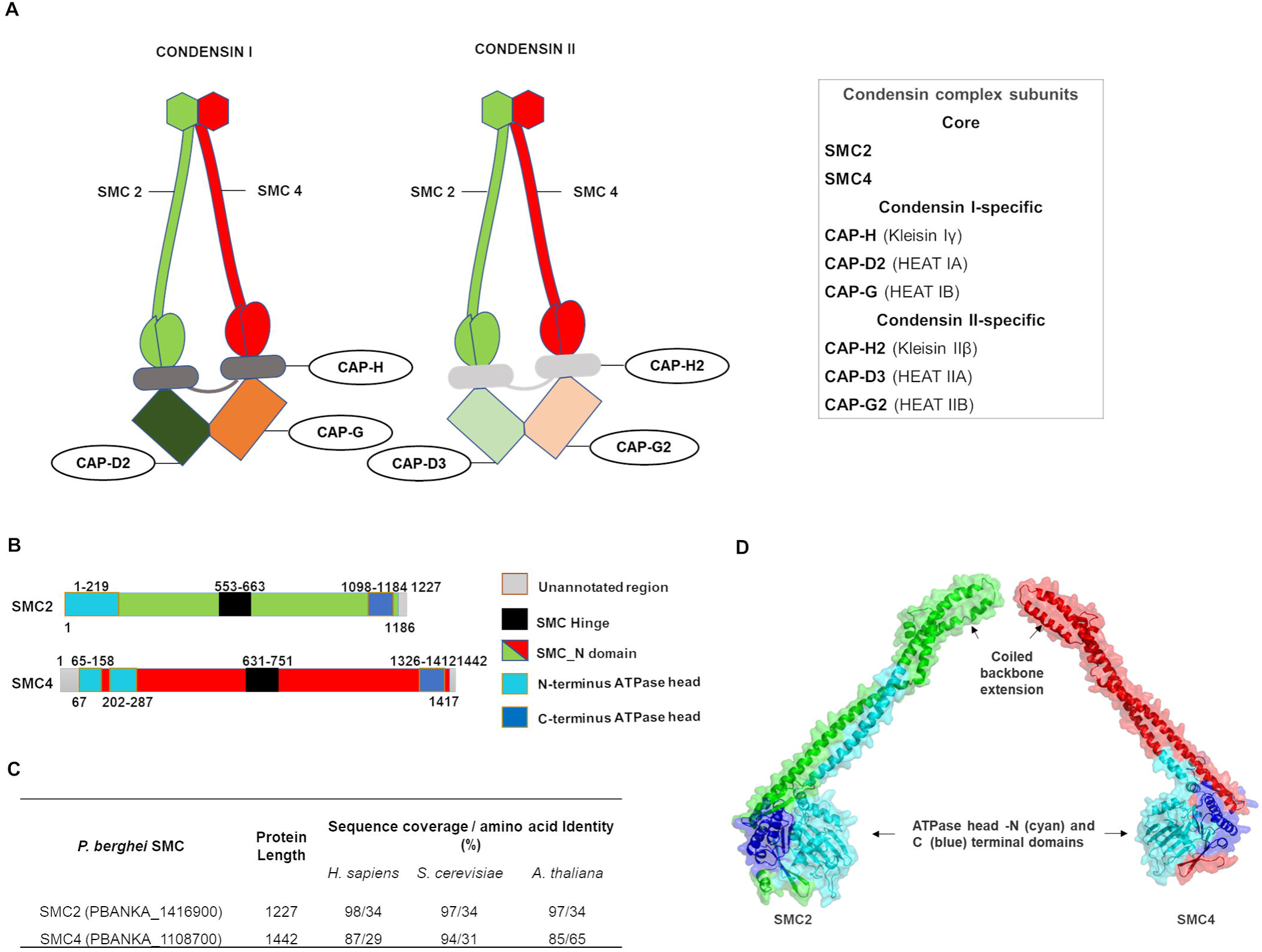
Architecture of condensin (SMC2/SMC4) in *Plasmodium berghei.* (A) Composition of two conventional condensin complexes (condensin I and condensin II) which are comprised of heterodimeric core subunits, SMC2 and SMC4, along with non-SMC regulatory subunits: Kleisin and a pair of HEAT subunits specific for either condensin I or II. Kleisin Iγ (CAP-H), Heat IA (CAP-D2) and Heat IB (CAP-G) form the condensin I complex, whereas Kleisin IIβ (CAP-H2), Heat IIA (CAP-D3) and Heat IIB (CAP-G2) form the condensin II complex (modified from Hirano T., 2016 and Uhlmann F., 2012). (B) Domain architecture of *P. berghei* SMC2 and SMC4. (C) Sequence coverage and amino acid identity of *P. berghei* SMC2 and SMC4 with *H. sapiens, S. cerevisiae* and *A. thaliana* proteins. (D) Homology-based predicted three-dimensional structures of *P. berghei* SMC2 and SMC4 showing coiled backbone extension without hinge domain and with ATPase head formation, required for condensin complex. See also Figure S1.

*Plasmodium*, the apicomplexan parasite that causes malaria, undergoes two types of atypical mitotic division during its life cycle: one in the asexual stages (schizogony in the liver and blood stages within the vertebrate host, and sporogony in the mosquito gut), and the other in male gametogenesis (endoreduplication) during the sexual stage (Arnot et al., 2011; Sinden, 1991b). Division during schizogony/sporogony resembles closed endomitosis with repeated asynchronous nuclear divisions, followed by a final synchronized set of nuclear division forming a multinucleated syncytium prior to cytokinesis. An intact nuclear envelope is maintained, wherein the microtubule organizing center (MTOC), known as the centriolar plaque or spindle pole body (SPB), is embedded, and rounds of mitosis and nuclear division proceed without chromosome condensation (Arnot et al., 2011; Francia and Striepen, 2014; Gerald et al., 2011; Sinden, 1991a, b; Sinden et al., 1976). In male gametogenesis, exposure of the male gametocyte to the mosquito midgut environment leads to activation of mitosis, which results in three rounds of rapid chromosome replication (endoreduplication) within 8-10 min and atypical chromosomal condensation, followed by nuclear and cell division to produce eight motile male gametes (exflagellation) (Guttery et al., 2012b; Sinden, 1991b; Sinden et al., 1976; Sinden et al., 2010). During exflagellation, each condensed haploid nucleus and associated MTOC, together with a basal body, axoneme and flagellum, form the microgamete that egresses from the main cellular body (Guttery et al., 2012b; Sinden, 1991b; Sinden et al., 1976; Sinden et al., 2010).

The atypical cell division and proliferation of malaria parasites is controlled by, amongst others, unique and divergent Apicomplexa-specific CDKs, aurora-like kinases (ARKs), mitotic protein phosphatase 1 (PP1), and only four APC components (Guttery et al., 2014; Roques et al., 2015; Wall et al., 2018; Ward et al., 2004; Wilkes and Doerig, 2008). However, there are no known classical group 1 cyclins, polo-like kinases (that are major regulators in mitotic entry) or classical mitotic protein phosphatases (CDC14 and CDC25) encoded in the genome (Guttery et al., 2014; Tewari et al., 2010; Ward et al., 2004; Wilkes and Doerig, 2008).

In *Plasmodium*, the role of condensin during cell division and general chromosome dynamics is unknown. Here, we investigated the location and function of the core subunits of condensin (SMC2 and SMC4) during two different mitotic division stages in the *Plasmodium* life cycle: during schizogony in the host’s blood, and during male gametogenesis in the mosquito vector. This study was performed using the rodent malaria model *Plasmodium berghei.* For this analysis we used a combination of cell biology, proteomics, transcriptomics, chromatin immunoprecipitation and reverse genetics approaches. Spatiotemporal localization using live cell imaging indicates a dynamic profile for both SMC2 and SMC4, with either discrete foci during early schizogony or a more diffuse nuclear localization during late schizogony and male gametogenesis. Genome-wide distribution studies using ChIP-seq experiments suggested that both components (SMC2/SMC4) are located at or near the centromeres during early schizogony, but this strong interaction was not observed during gametogenesis. Interestingly, we identified a differential composition of the condensin complex between the distinct mitotic stages, suggesting divergent mechanisms at the molecular level. Our data demonstrate that the condensin core subunits (SMC2/SMC4) have distinct functions at different stages of the parasite life cycle. Functional analyses using a conditional gene knockdown approach indicate that condensins are required for parasite proliferation and transmission.

## Results

### Bioinformatic analysis shows SMC2/SMC4, the core subunits of condensin, are encoded in the ***Plasmodium*** genome

In order to identify condensin in *Plasmodium* we screened for the core subunit genes in the *P. berghei* genome using PlasmoDB version 42, revealing both core SMC components of condensin, SMC2 and SMC4 (Bahl et al., 2002). Domain analysis revealed a conserved domain architecture for both SMC2 and SMC4 (**Figure 1B**). A comparative sequence analysis revealed low sequence similarity and identity (∼29-34%), except for the SMC4 homologue in *Arabidopsis thaliana* (65%) (**Figure 1C**), although there was similarity in size and overall domain structure when compared with the proteins in the other studied organisms. Interestingly, we found the *P. berghei* SMC4 N-terminal ATPase domain divided into two by a 44 amino acid insertion; a similar pattern has also been observed in other *Plasmodium* species. Subsequently, we generated a 3D model of the *P. berghei* SMC2 and SMC4 ATPase head domains and partial coiled region, using homology-based 3D structure modeling (**Figure 1D**). Root mean square deviation (RMSD) analysis of the 10 ns molecular dynamics (MD) simulation trajectory of the proteins showed a stable conformation comparable to pre-simulation energy-minimized structures. Radius of gyration analysis also confirmed a stable conformation for the predicted SMC2 and SMC4 domain structures during the 10 ns MD simulation (**Figure S1**). In this model of the SMC subunits, the N- and C-terminal ABC ATPase head and coiled-coil arms connecting the Hinge domain (**Figure 1D**) are present, as in other organisms. It is most likely that the heads of *Plasmodium* SMC2 and SMC4 undergo ATP-dependent engagement and disengagement, and they may perform chromosomal functions similar to those in other eukaryotes (Hirano, 2016).

### Condensin core subunits are expressed at every proliferative stage of the parasite life cycle and have a centromeric location during early schizogony

To locate the condensin SMC subunits during two proliferative stages (schizogony and male gametogenesis) of the *Plasmodium* life cycle, transgenic parasite lines were created to express GFP-tagged SMC2 and SMC4, using single crossover homologous recombination (**Figure S2A**). Integration PCR and western blot experiments were used to confirm the successful generation of transgenic lines (**Figure S2B and S2C**). We found that SMC2 and SMC4 are expressed during both schizogony and male gametogenesis. In early schizonts within host red blood cells, we observed discrete foci in the parasite cell adjacent to the nuclear DNA for both SMC2 and SMC4 whereas in mature schizonts the signal was more dispersed throughout the nucleus (**Figure 2A and 2B**). During male gametogenesis the proteins were also dispersed throughout the nucleus (**Figure 2C and 2D**). To further validate the SMC4 subcellular location, fractionation of cytoplasmic and nuclear extracts derived from purified gametocytes revealed the presence of SMC4 in the nucleus (**Figure S2D**). In addition, we also observed SMC4GFP distributed either as dispersed in the nucleus or at a discrete focus adjacent to the DNA throughout the parasite life cycle including in female gametocytes, in ookinetes, during oocyst development and in the liver stages (**Figure S3A-C**), suggesting that condensin core subunits are likely involved at all proliferative stages of the parasite life cycle. To examine whether these foci are centromeric or centrosomal, we used two approaches: we performed a co-localization experiment, using parasites expressing SMC4GFP crossed with those expressing NDC80mCherry, a kinetochore/centromeric marker protein (Cheeseman, 2014; McKinley and Cheeseman, 2016; Musacchio and Desai, 2017; Pandey et al., 2019), and we performed immunofluorescence assays using anti-centrin and anti-α-tubulin together with anti-GFP antibodies. Live imaging using the SMC4GFP and NDC80mCherry genetic cross showed co-localization of SMC4 and NDC80 in early schizonts and discrete foci of NDC80mCherry alone in gametocytes (**Figure 2E and 2F**). A similar pattern was observed in ookinetes and oocysts, with centromeric colocalization of SMC4 and NDC80 (**Figure S3D**). In early schizonts, immunofluorescence assays with anti-centrin antibodies revealed that SMC4 is located between centrin and the DAPI-stained nuclear DNA, confirming the non-centrosomal localization of SMC4 (**Figure 2G**). However, partial co-localization was observed with anti-α-tubulin antibodies in schizonts (**Figure 2G**) and during male gametogenesis (**Figure 2H**).

**Figure 2:**
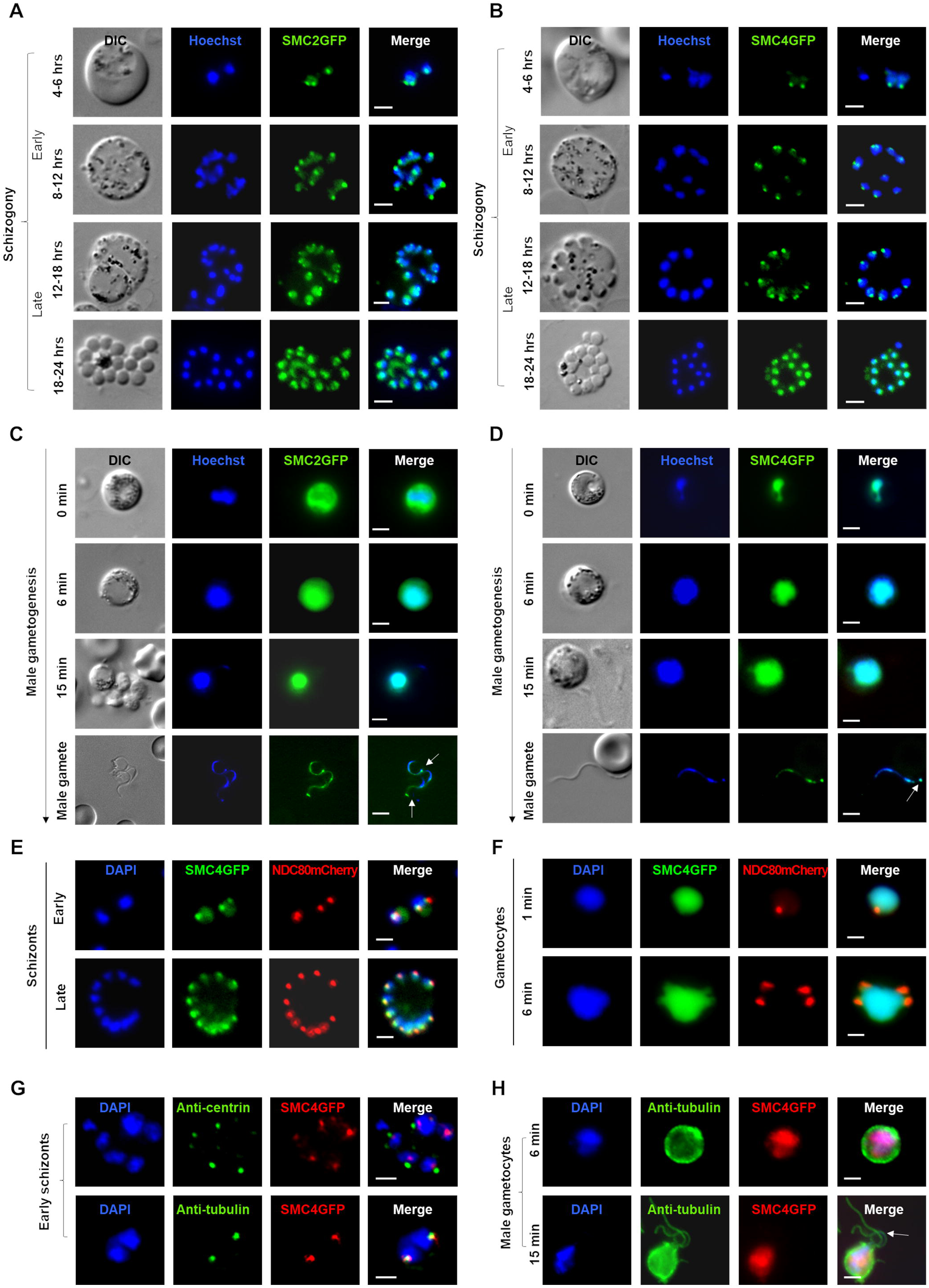
Temporal dynamics of condensin (SMC2 and SMC4) in two distinct *Plasmodium* proliferative stages (schizogony and male gametogenesis) undergoing atypical mitotic division. (A-D) Live cell imaging of SMC2GFP and SMC4GFP expressed during schizogony (100X magnification) and male gametogenesis (63X magnification). Time points indicate imaging done for schizont and gametocyte after the start of respective cultures. The white arrows in Panels C and D indicate discrete localization in male gametes. DIC: Differential interference contrast, Merge: Hoechst and GFP. Scale bar□= □2µm. (E-F). Live cell imaging of SMC4GFP and NDC80mCherry localization during schizogony and male gametogenesis. Merge: Hoechst, GFP and mCherry. Scale bar□= □2µm. (G-H) Immunofluorescence fixed-cell imaging of SMC4GFP and colocalization with antibodies specific for centrin and α-tubulin in mitotic cells (early schizonts 100X magnification and male gametocytes 63X magnification). The white arrow in Panel H indicates exflagellating male gamete. Scale bar□= □2µm. See also Figure S2, S3.

To identify the SMC2 and SMC4 DNA binding sites in a genome-wide manner, we performed ChIP-seq experiments for the schizont (after 8 hours in culture) and gametocyte (6 min post-activation) stages using SMC2GFP- and SMC4GFP-tagged parasites. A wild type strain (WTGFP) was used as a negative control. Binding of the SMC2 and SMC4 subunits was restricted to a region close to the previous computationally-annotated centromeres (centromere locations of *P. berghei* chromosomes had been predicted using *P. falciparum* as a reference and the conservation of genomic sequences among *Plasmodium* spp.) of all 14 chromosomes at the early schizont stage (**Figure 3A**) (Iwanaga et al., 2012). At this stage we observed significant ChiP-seq peaks with an average of 14.6- and 12.7-fold change (FC) compared to background in all pericentromeric regions for both SMC2 and SMC4 respectively. This restriction was not observed during gametogenesis, with instead a random distribution of condensin core subunits. While non-significant peaks (less than 1.5-FC) were detected in pericentromeric regions for SMC2, an even smaller increase in ChIP-Seq coverage in pericentromeric regions (1.08-FC) observed for SMC4 lead us to believe that the small peak observed during gametogenesis for SMC2 could be explained by a weak interaction of SMC2 or residual asexual signal in gametocyte samples. Identical patterns were obtained between biological replicates for each condition analyzed, confirming the reproducibility of the ChIP-seq experiments and suggesting a distinct function for the core subunits in these two mitotic stages. (**Figure 3A**). To confirm the location of the kinetochores/centromeres we also performed ChIP-seq with activated gametocytes from the NDC80GFP line (Pandey et al., 2019). Strong ChIP-seq peaks with an average of 74.8 FC were observed at the centromeres of all 14 chromosomes with perfect overlap with SMC2/SMC4 signals. These data clearly confirmed that the SMC2/SMC4 location during early schizogony was the centromeric location of NDC80 (**Figure3A**).

**Figure 3:**
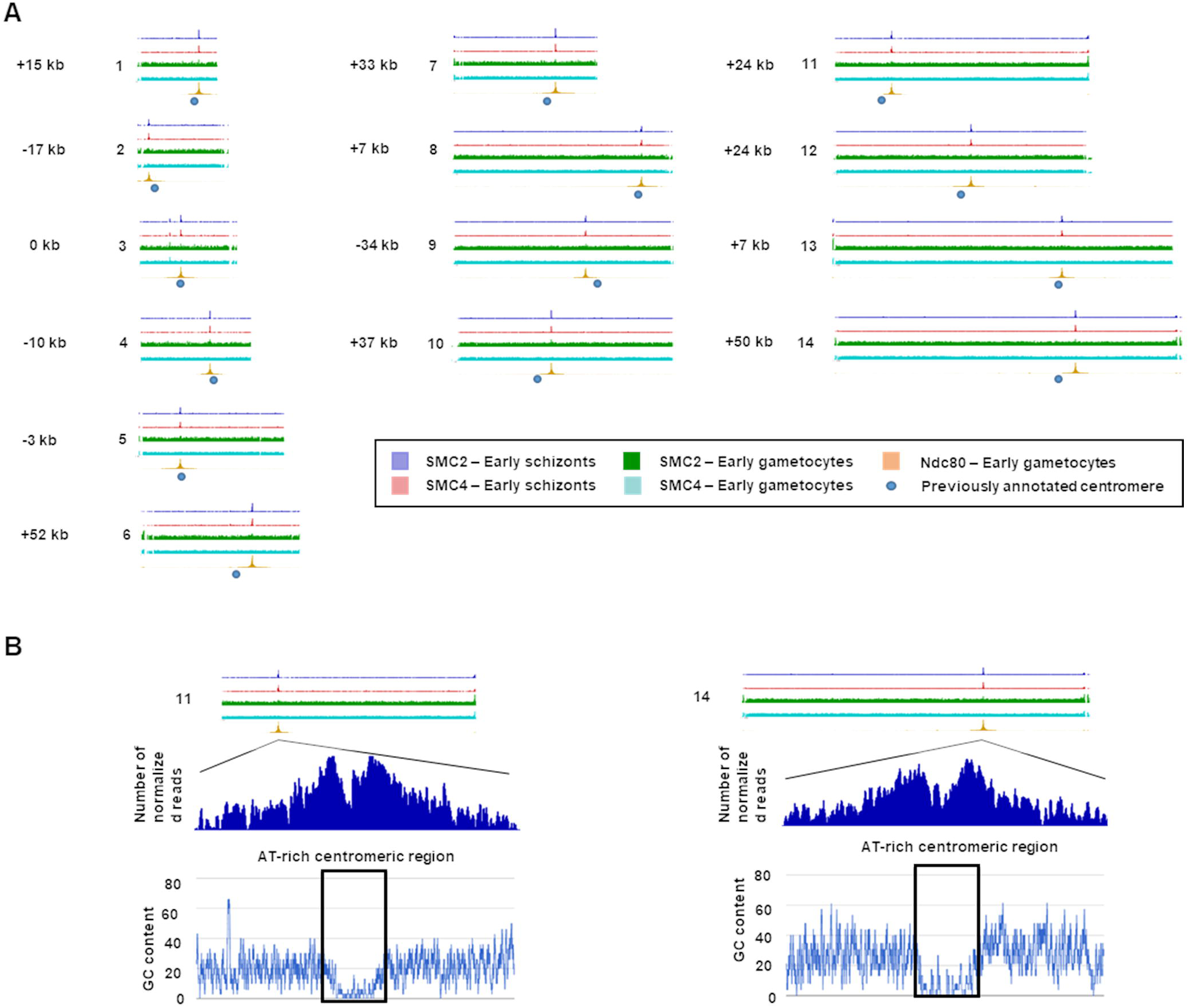
ChIP-seq analysis of SMC2GFP, SMC4GFP and NDC80 profiles. (A) Genome-wide ChIP-seq signal tracks for SMC2GFP and SMC4GFP for all 14 chromosomes in schizont and gametocyte stages. Both SMC schizont tracks and the NDC80 track each represent the average of two biological replicates, while the SMC4 gametocyte track represents the average of four biological replicates, one of which had two technical replicates averaged together. The location of previously annotated centromeres is indicated by blue circles. SMC2 and SMC4 proteins bind near the putative centromere in each of the 14 chromosomes (distance from centromere is shown in ±kb). Centromeric location was confirmed by genome-wide ChIP-seq signal tracks for NDC80GFP for all 14 chromosomes in gametocyte stage. Note that the scale for all SMC tracks is between 0-20 normalized read counts, while the NDC80 track is between 0-150 normalized read counts because of the considerable fold change enrichment observed for the NDC80 ChIP-seq signal. (B) Zoom-in on regions associated with the identified ChIP-seq peak. Low GC-content at the centers of peaks shown for chromosome 11 and chromosome 14 suggests association of the proteins with these newly defined centromeres in the schizont stage. Signals are plotted on a normalized read per million (RPM) basis. See also Table S1.

The chromosome binding sites detected for NDC80, and SMC2 and SMC4 were slightly offset from the locations previously annotated as centromeres (Iwanaga et al., 2012). However, the ChIP-seq peaks for all 14 chromosomes were centered on distinct regions of very low GC-content that are not present in the previously annotated centromeric regions (shown for chromosomes 11 and 14 in **Figure 3B**). AT-rich troughs have been associated with centromeres in yeasts (Lynch et al., 2010). The peaks are also located within extended intergenic regions indicating that the NDC80/SMC binding sites are the first experimentally validated centromeres for all 14 *P. berghei* chromosomes (**Table S1**).

### The full condensin complex is present during male gametogenesis and in mature schizonts, but ancillary proteins are absent in early schizonts

To examine the co-localization of SMC2 and SMC4 proteins, we generated transgenic parasite lines expressing either SMC2mCherry or SMC4GFP and crossed them genetically. The progeny, expressing both SMC2mCherry and SMC4GFP, showed co-localization of the two proteins during schizogony and gametogenesis (**Figure 4A**) consistent with SMC2 and SMC4 heterodimer complex formation at both stages.

**Figure 4:**
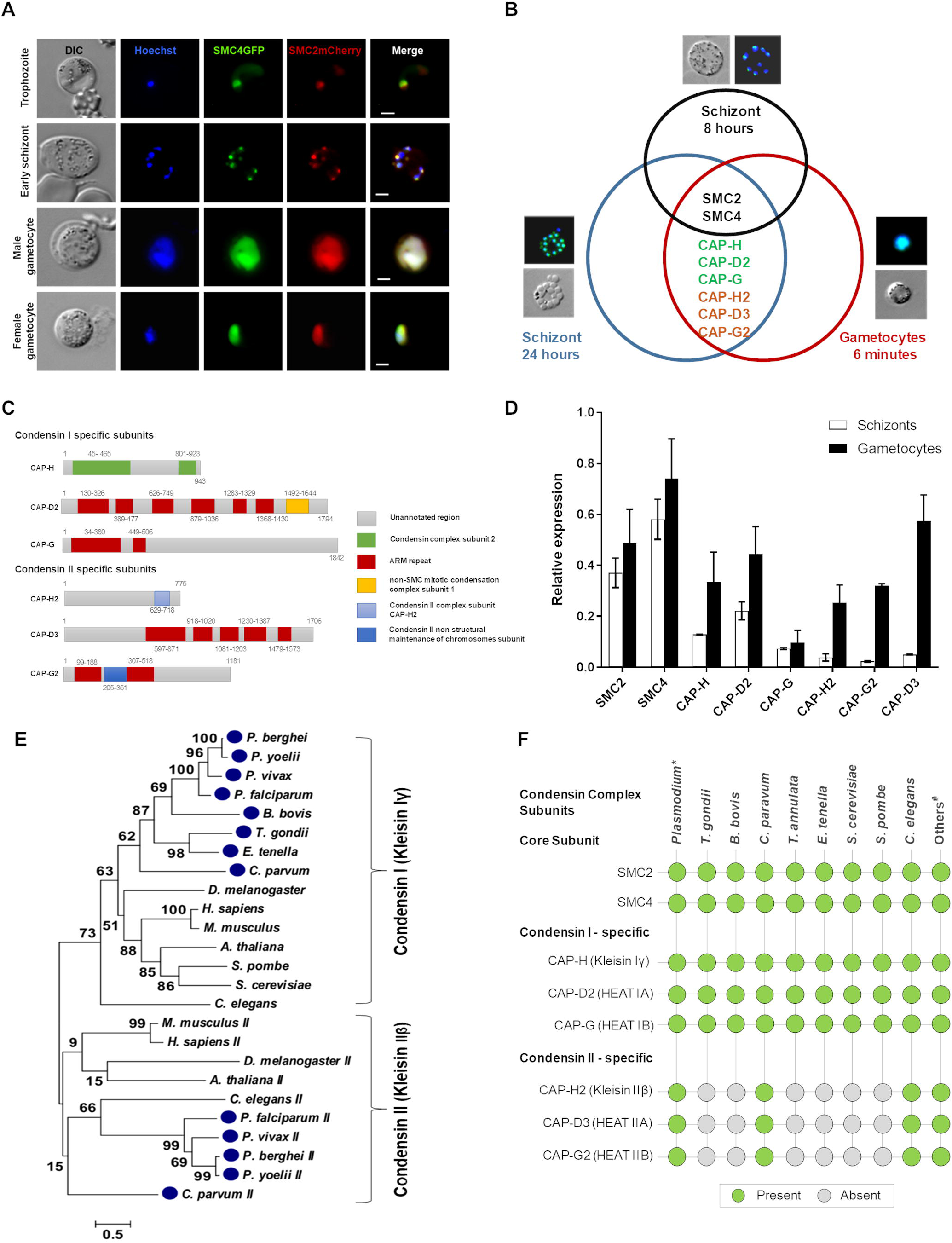
Differential condensin complex formation during schizogony and male gametogenesis, and phylogenetic analysis of kleisin. (A) Co-localization of SMC4GFP (green) and SMC2mCherry (red). Merge: Hoechst, GFP and mCherry. Scale bar□= □2µm. (B) Venn diagram displays the unique and shared proteins in the condensin complex of schizonts and gametocytes. Analysis of SMC2GFP and SMC4GFP protein complexes was done by tryptic digestion and liquid chromatography-mass spectrometry following GFP-specific immunoprecipitation from a lysate of schizonts maintained in culture for 8 hours and 24 hours, and gametocytes activated for 6 min. The representative live cell pictures have been taken from Figure 2. The list of all identified proteins is provided as Table S2. (C) Different domain architecture for subunits of Condensin I and Condensin II complexes. Schematic figure displays domain composition and protein length in the respective complex subunits. (D) qRT-PCR analysis of condensin complex subunit expression in schizont and gametocyte stages of parasite life cycle. Error bar = ±SD, n= 3. (E) Maximum likelihood phylogeny based on the alignment of kleisin subunits from apicomplexan species (*Plasmodium* spp, *Toxoplasma gondii, Cryptosporidium parvum, Babesia bovis* and *Eimeria tenella)* and other selected organisms. Topological support from bootstrapping is shown at the nodes. The protein sequences for selected organisms have been provided in File S1. (F) Distribution of condensin components across Apicomplexa and other organisms. Presence (green circle) or absence (grey circle) of condensin complex genes in each genome. * represent 4 *Plasmodium* spp., namely, *P. falciparum, P. vivax, P. berghei* and *P. yoelii*; # denotes *H. sapiens, A. thaliana* and *D. melanogaster*. See also Figure S4, Table S2 and File S1.

Next, we directly investigated the interaction between SMC2 and SMC4, and the presence of other potential interacting partner proteins, such as other condensin components (**Figure 1A**). We immunoprecipitated SMC2GFP and SMC4GFP from lysates of cells undergoing asexual endomitotic division at two time points (early schizogony, following 8 hours incubation in schizont culture medium *in vitro*, when most parasites are undergoing nuclear division and show discrete SMC2/4 foci, and after 24 hours incubation in schizont culture medium, when most parasites are mature schizonts or free merozoites with a dispersed SMC2/4 location). We also immunoprecipitated the proteins from parasites undergoing gametogenesis (at 6 min after activation, when the chromosomes are beginning to condense and cells are in the last phase before cytokinesis). Immunoprecipitated proteins were then digested with trypsin and the resultant peptides analyzed by liquid chromatography and mass spectrometry (LC-MS/MS). From all three samples, we recovered peptides from both SMC subunits confirming SMC2-SMC4 heterodimer formation during both schizogony and male gametogenesis (Figure 4B). From early schizonts, only SMC2- and SMC4-derived peptides were recovered, whereas from mature schizonts and gametocytes, we detected kleisins and other components of canonical condensin I and II complexes together with the SMC subunits except for CAP-G (**Figure 4B and Table S2**). In some early schizont samples, condensin II HEAT subunits (CAP-G2 and CAP-D3) were observed; however, kleisin was never recovered in five early schizont experimental replicates, and therefore we assume that formation of the complete condensin II complex does not occur at this stage (**Table S2**). All conventional condensin I and II complex subunits were identified by *in silico* analysis of the *Plasmodium* genome; for example, a BLAST search using fission yeast CAP-G revealed *Plasmodium* merozoite organizing protein (MOP) to be *Plasmodium* CAP-G. This is in agreement with the immunoprecipitation data in which we detected Kleisin Iγ (CAP-H), Heat IA (CAP-D2) and Heat IB (CAP-G, annotated as MOP) from the condensin I complex, and Kleisin IIβ (CAP-H2), Heat IIA (CAP-D3) and Heat IIB (CAP-G2) from the condensin II complex. The predicted domain architecture of these subunits from *P. berghei* is shown in **Figure 4C**. We also investigated the expression profile of condensin complex subunits in schizont and gametocyte stages of the parasite life cycle. We performed qRT-PCR for all eight components of the condensin complexes. The results revealed comparatively high expression of all condensin subunits in gametocytes as compared to schizonts (**Figure 4D**). In addition, levels of non-SMC condensin II components expressed in schizonts were much lower than those of non-SMC condensin I components, whereas in gametocytes comparable expression levels were observed for condensin I and II components except for CAP-G (**Figure 4D**). In ookinetes, the expression of non-SMC condensin components was low, except for CAP-H, which showed a level similar to that of the SMC components (**Figure S4**). Chromosome condensation in schizonts has not been reported, so why all components of both condensin complexes are present in mature schizonts is unclear. The presence of both condensin I and II in male gametocytes is consistent with the potentially atypical chromosomal condensation that has been previously observed in male gametocytes just before exflagellation (Sinden, 1991b; Sinden et al., 1976; Sinden and Hartley, 1985).

In view of the importance of kleisin to the structure and function of the condensin complexes, we examined the evolutionary relationships of kleisin among some apicomplexans and other organisms including *S. cerevisiae, A. thaliana* and *Homo sapiens*. The phylogenetic analysis indicated that kleisin is clustered into two groups, which correspond to components of condensin I and condensin II, respectively (**Figure 1A and 4E**). The presence of both condensin I and II component genes only in *Plasmodium* and *Cryptosporidium* shows that requirement for both condensin complexes is not a universal feature of Apicomplexa (**Figure 4F**). Other apicomplexans, for example *Toxoplasma* and *Eimeria*; have only condensin I components, similar to yeast homologues. These data suggest that *Plasmodium* and *Cryptosporidium* possess features of chromosome condensation and segregation that are distinct from those of other members of the phylum.

### Knockdown of condensin (SMC2 and SMC4) expression affects parasite proliferation and impairs parasite transmission

To examine further the functions of SMC2 and SMC4, we first attempted to delete the two genes. In both cases we were unable to produce gene knockout (KO) mutants (**Figure S5A**). Similar results have been reported previously from large-scale genetic screens in *P. berghei* (Bushell et al., 2017; Schwach et al., 2015). Together these data indicate that the condensin subunits SMC2 and SMC4 are likely essential for asexual blood stage development (schizogony). To investigate the function of SMC2 and SMC4 during cell division in male gametogenesis, we used a promoter trap double homologous recombination (PTD) approach to down-regulate gene expression at this stage by placing each of the two genes under the control of the AMA1 promoter (**Figure S5B**). AMA1 is known to be highly expressed in asexual blood stages but not during sexual differentiation. This strategy resulted in the successful generation of two transgenic parasite lines: *P*_*ama1*_*smc2* (SMC2PTD) and *P*_*ama1*_*smc4* (SMC4PTD), respectively (**Figure S5C**).

Since SMC2PTD and SMC4PTD had similar phenotypes, we performed a global transcriptome analysis only on SMC4PTD to identify other affected genes and regulators involved in cell division and proliferation. This analysis of SMC4PTD gametocytes 30 min after activation (when chromosome condensation and exflagellation are complete) confirmed the nearly complete ablation of *smc4* gene expression (**Figure 5A**). For pairs of the two biological samples (wild-type [WTGFP] and SMC4PTD), the Spearman correlation coefficients of 0.97 and 0.99 respectively, demonstrate the reproducibility of this experiment. (**Figure 5B**). In addition to SMC4, expression of a further 104 genes was also significantly downregulated, while expression of only 5 genes was significantly upregulated (**Figure 5C and Table S3**). Gene Ontology (GO) enrichment analysis of the downregulated genes identified several associated with microtubule and cytoskeletal function (**Figure 5D**). The reduced expression levels of ten of these genes was also examined by qRT-PCR (**Figure 5E**). By this method, there was a statistically significant difference in the level of expression of nine of these genes when comparing samples from WTGFP and SMC4PTD (**Figure 5E**). Of particular interest are AP2O2 (an AP2 domain transcription factor) and HMG (putative high mobility group protein B3), which act as transcription regulators in ookinetes; a putative SET domain protein which is known to be involved in methyl group transfer from S-adenosyl-L-methionine (AdoMet) to a lysine residue in histones and most likely associated with transcriptional repression; and finally an RCC, a protein predicted to be involved in chromosome condensation and chromosomal dynamics (Bahl et al., 2002). Other genes that were significantly downregulated include FRM2, involved in cytoskeleton organization; CCRNOT2 and NOT that form the CCR4-NOT complex, a key regulator of eukaryotic gene expression; and SEC7, involved in regulation of ARF protein signal transduction. Some of the other significantly downregulated genes, include AP2 transcription factor AP2-Sp; molecular motor kinesin-4, a putative regulator of chromosome condensation (PBANKA_0820800); and SMC1, a member of the SMC family, all known to be involved in either gene expression or chromatid segregation.

**Figure 5:**
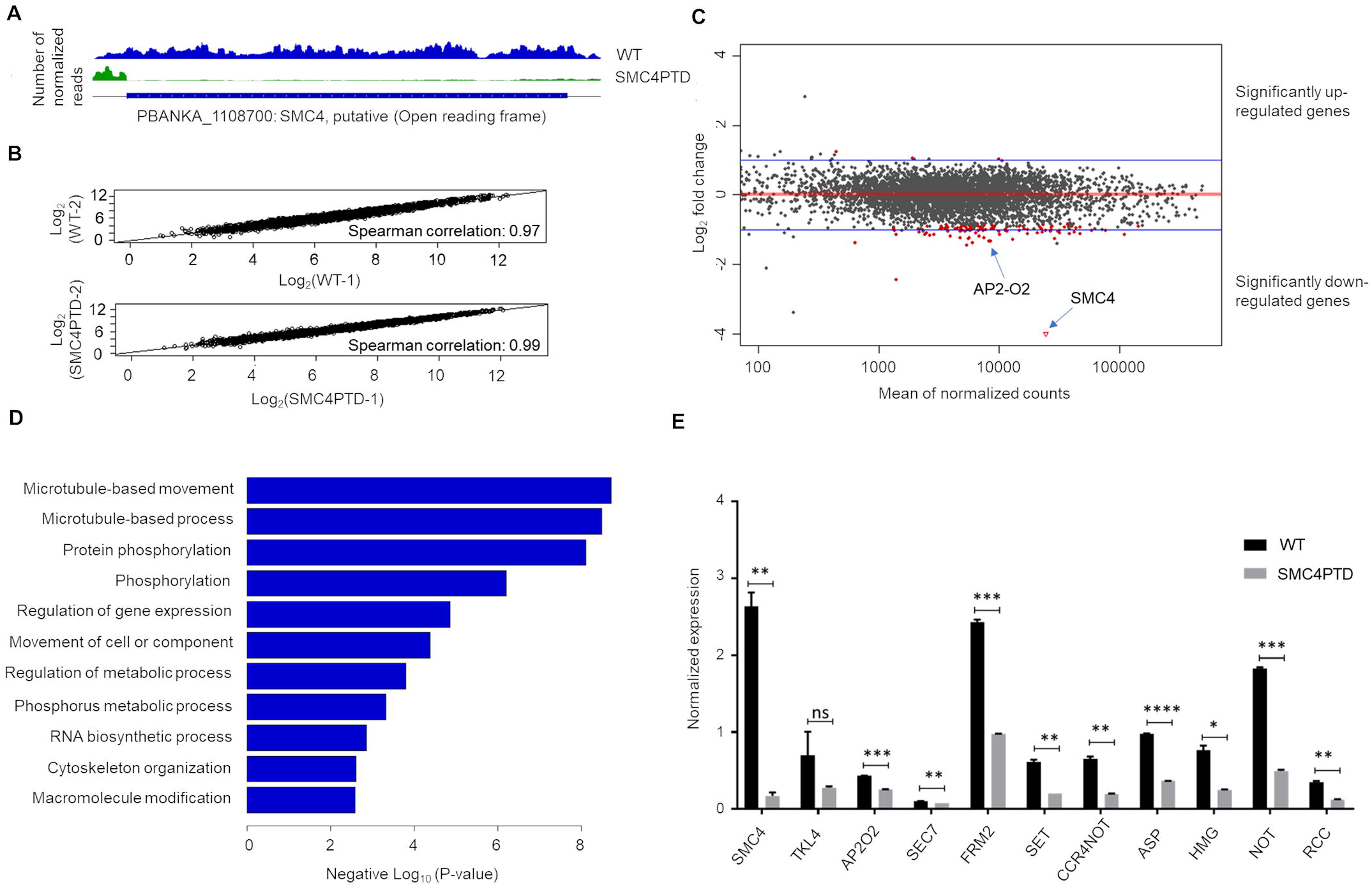
Global transcriptomic analysis for SMC4PTD in activated gametocytes by RNA-seq. (A) Confirmation of successful depletion of SMC4 transcript in the SMC4PTD line. Tracks shown each represent the average of two biological replicates. (B) Log-normalized scatterplots demonstrating high correlation between replicates for genome-wide expression. Spearman correlation was calculated using read counts normalized by number of mapped reads per million. (C) MA plot summarizing RNA-seq results. M: Log ratio and A: Mean average. Every gene is placed according to its log fold expression change in the SMC4PTD line as compared to the WT line (y-axis), and average expression level across replicates of both lines (x-axis). Red color indicates statistical significance of differential expression at the false positive threshold of 0.05. 105 genes are downregulated in the SMC4PTD line, and 5 genes are upregulated. (D) GO enrichment analysis of genes with log_10_ fold expression change of −0.5 or lower in the SMC4PTD line. (E) qRT-PCR analysis of selected genes identified as down-regulated in (C), comparing transcript levels in WT and SMC4PTD samples. Error bar = ±SEM, n= 3. Unpaired t-test was performed for statistical analysis: *p<0.05 **p<0.001, ***p<0.0001 and ****p<0.00001. See also Table S3, Table S5 and Figure S5.

While we were unable to detect any particular impaired phenotype in the SMC2PTD and SMC4PTD lines at the asexual blood stage (schizogony), and the parasite formed a similar number of schizonts and nuclei as compared to the WTGFP line (**Figure 6A**), we observed a ∼50% reduction in the number of exflagellation centers during male gametogenesis (**Figure 6B**). Fertilization and zygote formation leading to ookinete conversion was reduced to 10-15% as compared to the WTGFP line (**Figure 6C**). The ookinete motility assay showed normal movement of SMC4PTD ookinetes as compared with WT (Video S1 and S2). In the mosquito gut on days 9, 14 and 21 post-infection, we detected significantly fewer oocysts in the SMC2PTD and SMC4PTD lines (**Figure 6D and 6E**). Furthermore, the oocysts were considerably smaller compared to those of WTGFP (**Figure 6D and Figure 6F**) with unequal distribution and clusters of DNA in some of the oocysts at 14 and 21 days post-infection. No sporogony or endomitosis was observed within oocysts (**Figure 6G**). We were also unable to detect any sporozoites in the mosquito salivary glands (**Figure 6H**) and hence no parasite transmission from infected mosquitoes to mice was observed for either SMC2PTD or SMC4PTD parasite lines in bite-back experiments (Table S4), indicating that condensins are required for parasite transmission.

**Figure 6:**
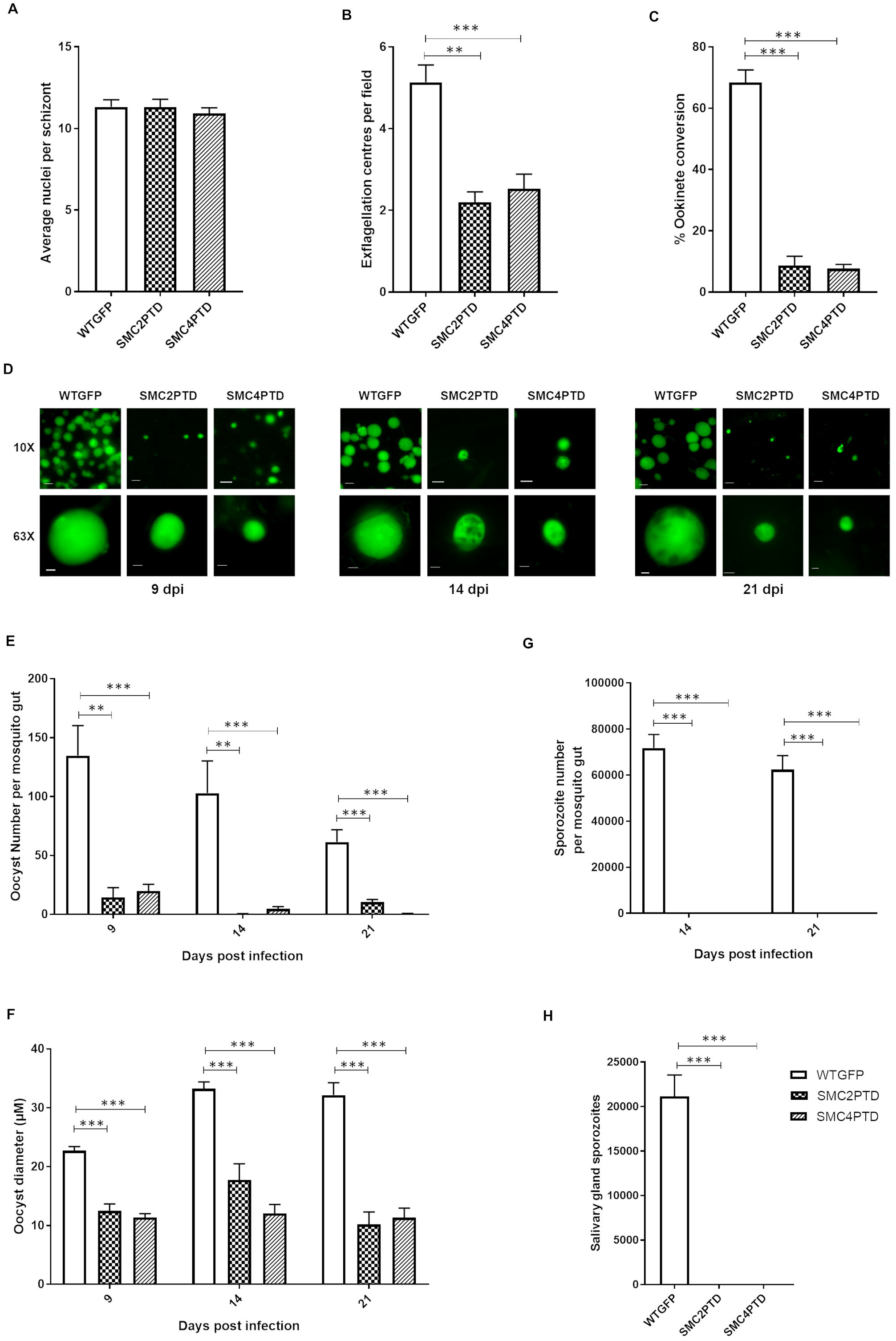
Phenotypic analysis of conditional gene expression knockdown in SMC2PTD and SMC4PTD transgenic lines at various proliferative stages during life cycle. (A) Average number of nuclei per schizont (mitotic division within red cell). N = 5 (minimum 100 cells). (B) Number of exflagellation centers (mitotic division during male gametogenesis) per field at 15 min post-activation. N = 3 independent experiments (10 fields per experiment) (C) Percentage ookinete conversion from zygotes. Minimum of 3 independent experiments (minimum 100 cells) (D) Live cell imaging of WTGFP, SMC2PTD and SMC4PTD oocysts (endomitosis in parasite within mosquito gut) at 9, 14 and 21 days post infection, using 10X and 63X magnification to illustrate differences in size and frequency. Scale bar□= □ 5 µm (63X) and 20 µm (10X). (E) Number of oocysts at 9, 14 and 21 days post-infection (dpi). N = 3 independent experiments with a minimum of 5 mosquito guts. (F) Oocyst diameter at 9, 14 and 21 days post-infection (dpi). N = 3 independent experiments. (G) Number of sporozoites at 14 and 21 dpi in mosquito gut. N = 3 independent experiments with a minimum of 5 mosquito guts. (H) Number of sporozoites at 21 dpi in mosquito salivary gland. Minimum of 3 independent experiments. Error bar = ±SEM. Unpaired t-test was performed for statistical analysis: *p<0.05 **p<0.01 and ***p<0.001. See also Table S4, Table S5, Figure S5 and Videos S1-S2.

## Discussion

Condensins are multi-subunit complexes that are involved in chromosomal condensation, organization, and segregation, and have been widely studied in many eukaryotes (Hirano, 2016; Uhlmann, 2016). The role of condensins in many unicellular protozoans such as *Plasmodium* remained elusive. Here we describe the structure, localization and functional role of the condensin core subunits (SMC2/SMC4) in the mouse malaria-causing parasite, *P. berghei* using molecular dynamics, live cell imaging, ChIPseq, protein pulldown and conditional gene knockdown approaches. *Plasmodium* shows atypical features of closed mitotic division resembling endomitosis during schizogony (with no observed chromosomal condensation and extensive asynchronous nuclear division followed by final round of synchronous nuclear division prior to cytokinesis), and endoreduplication in male gametogenesis (with rapid chromosome replication and atypical condensation prior to nuclear division, cytokinesis and exflagellation) (Arnot et al., 2011; Sinden, 1991b). Our previous studies have identified an unusual repertoire of proteins involved in the regulation of the parasite cell cycle and cell proliferation: there is no identifiable centrosome, no obvious complement of cell-cycle cyclins, a small subset of APC components, a set of divergent and *Plasmodium-*specific CDKs, and a complete absence of polo-like kinases and CDC24 and CDC14 phosphatases, as compared to most organisms that have been studied (Arnot et al., 2011; Francia et al., 2015; Guttery et al., 2012a; Guttery et al., 2014; Roques et al., 2015; Tewari et al., 2010).

Here, using bioinformatics screening, we showed that both condensin I and II complex subunit components are encoded in the *P. berghei* genome, as has been also described for *P. falciparum* in PlasmoDB (Bahl et al., 2002). The two core subunits of condensin, SMC2 and SMC4, have low sequence similarity to the proteins in model organisms but a similar protein structure as predicted by molecular modelling (Kelley et al., 2015).

Protein localization studies at different stages of the parasite life cycle using live cell imaging of SMC2GFP and SMC4GFP and immunofluorescence, show distinct patterns during the mitotic divisions of early and late schizogony and male gametogenesis. Whereas discrete protein foci were detected during endomitosis in early schizogony, a stage characterized by asynchronous nuclear division, a more dispersed nuclear localization was observed during late schizogony and male gametogenesis. By immunofluorescence, the discrete foci of SMC2/4GFP in early schizogony were located close to the stained DNA and close to but not coincident with centrin, marking the Spindle Pole Body (SPB). The ChIP-seq analyses suggest that in early schizonts SMC2 and SMC4 form a complex that binds at or near the centromere of all 14 chromosomes, a result that is further substantiated by the dual labelling and colocalization studies with the kinetochore/centromere marker, NDC80.The ChIP-seq analysis of NDC80GFP binding during gametogenesis also confirms the centromeric location of SMC2/SMC4. These results suggest that the SMC2/SMC4 complex alone is restricted to binding centromeric regions in the highly proliferative early schizont stage, where it may have a constrained role in sister chromatid cohesion and segregation (Iwasaki and Noma, 2016). Genome-wide studies of condensin distribution in mammalian or yeast cells have shown that the complex is non-randomly distributed across the chromosomes, and often found at the boundaries of topologically associating domains (TADs) within chromosome territories, which supports the proposed role in transcriptional regulation and global chromosomal organization (Kim et al., 2016; Yuen et al., 2017). As the *Plasmodium* genome lacks classical TADs (Ay et al., 2014; Bunnik et al., 2018; Bunnik et al., 2019) a restricted distribution of the condensin complex on the centromere of all 14 chromosomes suggests a distinct function.

Whilst we detected only the SMC2-SMC4 heterodimer in early schizogony, located within the nucleus at the centromere and at a discrete focus adjacent to but distinct from the SPB, the protein interaction analysis using SMC2/4-GFP showed that other subunits of the full condensin I and II complexes were present during late schizogony and male gametogenesis. It is thought that in late schizogony the last set of divisions is synchronous and followed by cytokinesis to produce mature merozoites and at this stage the dispersed distribution of SMC2/4-GFP was observed. A similar dispersed pattern of protein in the nucleus was observed during male gametogenesis, which is also associated with the presence of condensin complex I and II proteins preceding exflagellation. No chromosomal condensation has been reported in mature schizonts, although it has been observed in male gametogenesis, as shown by electron microscopy studies (Sinden, 1991b; Sinden et al., 1976; Sinden and Hartley, 1985). The presence of both condensin I and II complex in late schizogony suggests that the full complexes are only involved in the final synchronous cycle of nuclear division preceding cytokinesis. Previous studies have reported that non-SMC condensin II subunits are dispensable during schizogony but non-SMC components of condensin I complex are not (Bahl et al., 2002; Schwach et al., 2015). One of the components, CAP-G, also annotated as merozoite organizing protein (MOP), has been demonstrated to be essential for cytokinesis in *P. falciparum* asexual blood stages (Absalon et al., 2016). Our bioinformatics analysis shows that *Plasmodium* (two hosts, asynchronous cell division) and *C. parvum* (single host, with long duration dormant phase outside of host) are the only two Apicomplexan parasites that have components of both condensin I and II complexes encoded in the genome, similar to what is observed in higher eukaryotes. They also have unusual modes of cell division as compared with others such as *Toxoplasma* and *Babesia*, which display symmetrical modes of division (Francia and Striepen, 2014). Apicomplexan parasites that encode only a single condensin complex show no chromosome condensation. Similarly, other parasites, with closed mitosis and no chromosome condensation, for example *Trypanosoma brucei*, encode only condensin I complex (Hammarton, 2007), whereas in parasites with chromosome condensation, such as *Giardia intestinalis*, both condensin I and condensin II are present. However, Giardia lacks one of the conventional non-SMC HEAT subunits (HEAT IB and HEAT IIB) whereas HEAT IA and HEAT IIA are present (Tumova et al., 2015). Another protist, *Tetrahymena thermophila*, shows chromosomal condensation, exhibits noncanonical division between somatic and germline cells, and has an expanded set of condensin I paralogues, with different kleisin components between germline (Cph1 and Cph2) and somatic cells (Cph3, Cph4 and Cph5) (Howard-Till and Loidl, 2018; Howard-Till et al., 2019).

Condensin I and II complexes display distinct localization patterns in various organisms. In the red alga *Cyanidioschyzon merolae* (Fujiwara et al., 2013), condensin II has a centromeric location during metaphase, whereas condensin I distributes more broadly along the chromosome arms. In higher eukaryotes including *Drosophila melanogaster* (Oliveira et al., 2007), *Caenorhabditis elegans* (Collette et al., 2011) and HeLa cells (Hirota et al., 2004; Ono et al., 2004) condensin I is present in the cytoplasm and has a chromosomal location after the nuclear envelope is dissolved in open mitosis. The nuclear localization of condensin II is observed in interphase, it is stabilized on chromatin during prophase, and the complex remains associated with chromosomes throughout mitosis, at least in HeLa cells. Budding yeast and fission yeast, which undergo closed mitosis like *Plasmodium*, have only a single condensin complex, but there is a differential pattern of subcellular location in each species. In budding yeast, the condensin I complex is located in the nucleus throughout the cell cycle, a pattern observed for condensin II in higher eukaryotes, despite the greater protein sequence similarities of the yeast complex to higher eukaryote condensin I (Thadani et al., 2012). Also, within the nucleus, the condensin location at the kinetochore is cell cycle dependent (Bachellier-Bassi et al., 2008). In fission yeast, the single condensin complex is predominantly cytoplasmic during interphase and nuclear during mitosis, with the location dependent on CDK phosphorylation at Thr19 of SMC4/Cut3 (Sutani et al., 1999).

The present study shows that the SMC2-SMC4 complex plays an essential role during schizogony, as we, and previous genome-wide functional screens (Bushell et al., 2017), were unable to disrupt the genes. Our conditional knockdown using the promoter trap (PTD) approach suggests that reduction of both SMC2 and SMC4 expression affects both male gametogenesis and zygote differentiation, and causes total impairment of endomitotic cell division in the oocyst, thereby blocking parasite transmission.

The partial defect observed in male gamete formation (exflagellation) in the PTD parasite lines may be due to the necessity of condensin complex formation for proper chromosomal condensation during exflagellation. Transcriptomic analysis of the SMC4PTD line confirmed the reduced expression of the *smc4* gene and identified dysregulated transcripts that are likely either critical for gene expression, or chromosomal segregation and condensation, microtubule assembly and male gametocyte activation. This further demonstrates that SMC2 and SMC4 complexes are essential for proper chromosome condensation and separation during exflagellation. Among the significantly dysregulated genes, deletion of AP2-O2 has been shown to strongly impair ookinete and oocyst development, leading to an absence of sporozoite formation and a complete blockage of transmission (Modrzynska et al., 2017). The SET protein, which is a post translational modification protein, is essential for parasite survival (Schwach et al., 2015). RCC is predicted to be a regulator of chromosome condensation, is essential for parasite survival, and acts as anchor for both parasite kinase (CDPK7) and phosphatase (PP1) (Lenne et al., 2018). The phenotype observed in SMC4PTD parasites may therefore reflect contributions from all these differentially regulated genes.

The reduction in mature ookinete formation in the SMC4PTD line, suggests an important role for condensin during meiosis as well. During this stage, chromosomal condensation has been observed (Sinden and Hartley, 1985), and this may be similar to the situation in *Arabidopsis*, where condensin is important in chromosomal condensation during meiosis (Smith et al., 2014). A severe defect in number and size of oocyst formation in the mosquito gut was also observed, and at this stage multiple rounds of endomitotic division give rise to thousands of sporozoites. The process requires ten or more rounds of DNA replication, segregation, and mitotic division to create a syncytial cell (sporoblast) with thousands of nuclei over a period of several days (Francia and Striepen, 2014; Gerald et al., 2011). The proper segregation of nuclei into individual sporoblasts is organized by putative MTOCs (Roques et al., 2019; Sinden and Strong, 1978). As condensin has been shown to play an important role in organizing MTOCs (Kim et al., 2014) it may be that in the absence of condensin the endomitotic division is impaired and no sporozoites are formed.. Many mutants reported to cause a defect in oocyst maturation, such as *Pb*MISFIT, PbCYC3, PPM5, kinesin-8X and G actin-sequestering protein (Bushell et al., 2009; Guttery et al., 2014; Hliscs et al., 2010; Roques et al., 2015; Zeeshan et al., 2019) did not cause a significant change in the *smc4* expression profile, nor was there a significant change in the expression of these genes in the present study, suggesting that the SMC4PTD mutant parasite defect in the oocyst is independent of *Pb*MISFIT, *Pb*CYC3 and *Pb*PPM5 function.

In summary, the present study shows that the condensin core subunits SMC2 and SMC4 play crucial roles in the atypical mitosis of the *Plasmodium* life cycle and may perform distinct functions during different proliferative stages; specifically, during early schizogony, the final chromosome segregation in the last nuclear division during late schizogony and chromosome condensation before nuclear division and exflagellation during male gametogenesis. Their removal or depletion causes impaired parasite development and blocks transmission. Additional analyses of the non-SMC components of condensin I and II will provide further insight into the function of condensin during *Plasmodium* cell proliferation.

## Acknowledgements

We thank Prof. Frank Uhlmann, The Francis Crick Institute, for stimulating discussions and advice on condensins, Dr Cleidiane Zampronio for assisting in mass spectrometry analysis and Julie Rodgers for insectary assistance. This project was funded by MRC project grants and MRC Investigators grants awarded to RT (G0900109, G0900278, MR/K011782/1) and BBSRC (BB/N017609/1). RP was supported by MRC grant MR/K011782/1. AAH was supported by the Francis Crick Institute, which receives its core funding from Cancer Research UK (FC001097), the UK Medical Research Council (FC001097), and the Wellcome Trust (FC001097). DG was supported by the Department of Biotechnology (DBT), Government of India, grant BT/BI/25/066/2012, KGLR was supported by the National Institutes of Allergy and Infectious Diseases and the National Institutes of Health (grants R01 AI06775 and R01 AI136511) and the University of California, Riverside (NIFA-Hatch-225935).

## Author contribution

RT, AAH and KGLR conceived and designed all experiments. RT, RP, SA, MB, RJW, MZ, ER, AF, DB, ED and SW performed the GFP tagging and conditional knockdown experiments; RRS performed liver stage imaging; RP, MZ, ED and RT performed protein pulldown experiments; ARB performed the mass spectrometry; RP and DG performed the phylogenetic analysis and molecular dynamics; SA, RP XML, GB, TH, KGLR and RT performed the RNA-seq and ChIP-Seq experiments; RP, SA, AAH, KGLR and RT analyzed the data; RP, SA, AAH, KGLR and RT wrote the manuscript and all others contributed to it.

## STAR* METHODS

### Lead Contact and Materials Availability

Further information and requests for resources and reagents should be directed to and will be fulfilled by the Lead Contact, Rita Tewari

### EXPERIMENTAL MODEL AND SUBJECT DETAILS

*P. berghei* ANKA line 2.34 (for GFP-tagging) or ANKA line 507cl1 (for gene deletion and promoter swap) parasites were used for transgenic line creation as described previously (Wall et al., 2018). All animal work done at the University of Nottingham has passed an ethical review process and has been approved by the United Kingdom Home Office. The work was carried out under UK Home Office Project Licenses (40/3344,30/3248 and PDD2D5182). Six to eight-week-old female Tuck-Ordinary (TO) (Harlan) or CD1 outbred mice (Charles River) were used for all experiments performed in UK.

Infections of mice in Bern (Switzerland) were performed in accordance with the guidelines of the Swiss Tierschutzgesetz (TSchG; Animal Rights Laws) and approved by the ethical committee of the University of Bern (Permit Number: BE132/16). Female Balb/c mice (6-8 weeks; Janvier laboratories, France) were used to maintain transfected parasites and for feeding of mosquitoes with parasites.

Mice were injected via an intraperitoneal or intravenous route. When parasitemia reached 2-5%, mice were euthanized in a CO_2_ chamber and parasites isolated following exsanguination. For feeding of mosquitoes, upon reaching a parasitemia of 7-15%, mice were anaesthetized with a terminal dose of ketamine:xylazine and when no longer reacting to touch stimulus were placed on a cage of approximately 150 mosquitoes.

## METHOD DETAILS

### Generation of transgenic parasites

GFP-tagged vectors were designed using the p277 plasmid vector and transfected as described previously (Guttery et al., 2014). Targeted gene deletion vectors were designed using the pBS-DHFR plasmid (Tewari et al., 2010). Conditional gene knockdown constructs (SMC2PTD and SMC4PTD) were designed using *P*_*ama1*_ (*pSS368*) (Sebastian et al., 2012). *P. berghei* ANKA line 2.34 (for GFP-tagging) or ANKA line 507cl1 (for gene deletion and promoter swap) parasites were transfected by electroporation as described previously (Wall et al., 2018). Genotypic analysis was performed using diagnostic PCR reaction and Western blot. All of the oligonucleotides used to confirm genetically modified tag and mutant parasite lines can be found in Table S5. For western blotting, purified schizonts were lysed using lysis buffer (10 mM TrisHCl pH 7.5, 150 mM NaCl, 0.5 mM EDTA and 1% NP-40). The lysed samples were boiled for 10 min at 95 °C after adding Laemmli buffer. The samples were centrifuged at maximum speed (13000g) for 5 min. The samples were electrophoresed on a 4–12% SDS-polyacrylamide gel. Subsequently, resolved proteins were transferred to nitrocellulose membrane (Amersham Biosciences). Immunoblotting experiment was performed using the Western Breeze Chemiluminescence Anti-Rabbit kit (Invitrogen) and anti-GFP polyclonal antibody (Invitrogen) at a dilution of 1:1250, according to the manufacturer’s instructions.

### Phenotypic Analysis and Live Cell Imaging

Phenotypic analyses of the transgenic parasite lines were performed at different points of parasite life cycle as described previously (Guttery et al., 2014). Briefly, infected blood was used to analyse asexual blood stages and gametocytes. Schizont culture was used to analyse different stages of asexual development. *In vitro* cultures were prepared to analyse activated gametocyte, exflagellation, zygote formation and ookinete development. For *in vitro* exflagellation studies, gametocyte-infected blood was obtained from the tails of infected mice using a heparinised pipette tip. Gametocyte activation was performed by mixing 100 µl of ookinete culture medium (RPMI 1640 containing 25□mM HEPES, 20% fetal bovine serum, 10□mM sodium bicarbonate, 50□µM xanthurenic acid at pH 7.6) with the gametocyte infected blood. Microgametogenesis was monitored at two time points to study mitotic division (6 and 15□min post activation [mpa]). For mosquito transmission and bite back experiments triplicate sets of 40-50 *Anopheles stephensi* mosquitoes were used. The mosquito guts were analysed on different days post infection (dpi); 9 dpi, 14 dpi and 21 dpi to check oocyst development and sporozoite formation. For live cell imaging, parasites were stained with Hoechst 33342 DNA stain before mounting for fluorescent microscopy. For immunofluorescence assay (IFA), the material was fixed using 2% and 4% paraformaldehyde (PFA) in microtubule stabilising buffer (MTSB:10 mM MES, 150 mM NaCl, 5 mM EGTA, 5 mM MgCl_2_, 5 mM glucose) for schizonts and gametocytes, respectively. Immunochemistry was performed using primary antibodies; anti-GFP rabbit antibody (Invitrogen) at 1:250 dilution, anti-alpha-tubulin mouse antibody (Sigma-Aldrich) at 1:1000 dilution, and anti-centrin mouse clone 20h5 antibody (Millipore) at 1:200 dilution. Secondary antibodies were AlexaFluor 568 labelled anti-rabbit (red) and AlexaFluor 488 labelled anti-mouse (green) (Invitrogen) (1:1000 dilution). The slides were mounted in Vectashield with DAPI (Vector Labs) for fluorescent microscopy. Parasites were visualised on a Zeiss AxioImager M2 microscope fitted with an AxioCam ICc1 digital camera (Carl Zeiss, Inc).

For liver stages, 1 × 10^5^ HeLa cells were seeded in glass-bottomed imaging dishes. HeLa cells were grown in MEM (minimum essential medium) with Earle’s salts, supplemented with 10% heat inactivated FCS (foetal calf serum), 1% penicillin/streptomycin and 1% l-glutamine (PAA Laboratories) in a humid incubator at 37°C with 5% CO_2_. 24 hours after seeding, sporozoites were isolated from parasite-infected mosquito salivary glands and used to infect seeded HeLa cells. Infected cells were maintained in 5% CO_2_ at 37°C. To perform live cell imaging, Hoechst 33342 (Molecular Probes) was added (1 μg/ml) and imaging was done at 48 h and 55 h post-infection using a Leica TCS SP8 confocal microscope with the HC PL APO 63x/1.40 oil objective and the Leica Application Suite X software.

### ChIP-seq and global transcriptomic analysis

For the ChIP-seq analysis, libraries were prepared from crosslinked cells (using 1% formaldehyde). The crosslinked parasite pellets were resuspended in 1 mL of nuclear extraction buffer (10 mM HEPES, 10 mM KCl, 0.1 mM EDTA, 0.1 mM EGTA, 1 mM DTT, 0.5 mM AEBSF, 1X protease inhibitor tablet), post 30 min incubation on ice, 0.25% Igepal-CA-630 was added and homogenized by passing through a 26G x ½ needle. The nuclear pellet extracted through 5000 rpm centrifugation, was resuspended in 130 µl of shearing buffer (0.1% SDS, 1 mM EDTA, 10 mM Tris-HCl pH 7.5, 1X protease inhibitor tablet), and transferred to a 130 µl Covaris sonication microtube. The sample was then sonicated using a Covaris S220 Ultrasonicator for 10 min for schizont samples and 6 min for gametocyte samples (Duty cycle: 5%, Intensity peak power: 140, Cycles per burst: 200, Bath temperature: 6°C). The sample were transferred to ChIP dilution buffer (30 mM Tris-HCl pH 8, 3 mM EDTA, 0.1% SDS, 30 mM NaCl, 1.8% Triton X-100, 1X protease inhibitor tablet, 1X phosphatase inhibitor tablet) and centrifuged for 10 min at 13,000 rpm at 4°C, retaining the supernatant. For each sample, 13 μl of protein A agarose/salmon sperm DNA beads were washed three times with 500 µl ChIP dilution buffer (without inhibitors) by centrifuging for 1 min at 1000 rpm at room temperature, then buffer was removed. For pre-clearing, the diluted chromatin samples were added to the beads and incubated for 1 hour at 4°C with rotation, then pelleted by centrifugation for 1 min at 1000 rpm. Supernatant was removed into a LoBind tube carefully so as not to remove any beads and 2 µg of anti-GFP antibody (ab290, anti-rabbit) were added to the sample and incubated overnight at 4°C with rotation. Per sample, 25 µl of protein A agarose/salmon sperm DNA beads were washed with ChIP dilution buffer (no inhibitors), blocked with 1 mg/mL BSA for 1 hour at 4°C, then washed three more times with buffer. 25 µl of washed and blocked beads were added to the sample and incubated for 1 hour at 4°C with continuous mixing to collect the antibody/protein complex. Beads were pelleted by centrifugation for 1 min at 1000 rpm at 4°C. The bead/antibody/protein complex was then washed with rotation using 1 mL of each buffers twice; low salt immune complex wash buffer (1% SDS, 1% Triton X-100, 2 mM EDTA, 20 mM Tris-HCl pH 8, 150 mM NaCl), high salt immune complex wash buffer (1% SDS, 1% Triton X-100, 2 mM EDTA, 20 mM Tris-HCl pH 8, 500 mM NaCl), high salt immune complex wash buffer (1% SDS, 1% Triton X-100, 2 mM EDTA, 20 mM Tris-HCl pH 8, 500 mM NaCl), TE wash buffer (10 mM Tris-HCl pH 8, 1 mM EDTA) and eluted from antibody by adding 250 μl of freshly prepared elution buffer (1% SDS, 0.1 M sodium bicarbonate). We added 5 M NaCl to the elution and cross-linking was reversed by heating at 45°C overnight followed by addition of 15 μl of 20 mg/mL RNAase A with 30 min incubation at 37°C. After this, 10 μl 0.5 M EDTA, 20 μl 1 M Tris-HCl pH 7.5, and 2 μl 20 mg/mL proteinase K were added to the elution and incubated for 2 hours at 45°C. DNA was recovered by phenol/chloroform extraction and ethanol precipitation, using a phenol/chloroform/isoamyl alcohol (25:24:1) mixture twice and chloroform once, then adding 1/10 volume of 3 M sodium acetate pH 5.2, 2 volumes of 100% ethanol, and 1/1000 volume of 20 mg/mL glycogen. Precipitation was allowed to occur overnight at −20°C. Samples were centrifuged at 13,000 rpm for 30 min at 4°C, then washed with fresh 80% ethanol, and centrifuged again for 15 min with the same settings. Pellet was air-dried and resuspended in 50 μl nuclease-free water. DNA was purified using Agencourt AMPure XP beads.

For the global transcriptome analysis, total RNA was isolated from parasite pellet using RNA extraction Kit and lyophilized. The NEB poly(A) mRNA magnetic isolation module (E7490L) was used to isolate mRNA, while the NEBNext Ultra Directional RNA Library Prep Kit (E7420L) was used to prepare a cDNA library from the isolated mRNA using manufacturer’s instructions.

Libraries were prepared using the KAPA Library Preparation Kit (KAPA Biosystems), and were amplified for a total of 12 PCR cycles (15 s at 98°C, 30 s at 55°C, 30 s at 62°C) using the KAPA HiFi HotStart Ready Mix (KAPA Biosystems). Libraries were sequenced using a NextSeq500 DNA sequencer (Illumina), producing paired-end 75-bp reads.

### Immunoprecipitation and Mass Spectrometry

Schizonts, following 8 hours and 24 hours respectively in *in vitro* culture, and male gametocytes 6 min post activation were used to prepare cell lysates. Purified parasite pellets were crosslinked using formaldehyde (10 min incubation with 1% formaldehyde, followed by 5 min incubation in 0.125M glycine solution and 3 washes with phosphate buffered saline (PBS) pH7.5). Immunoprecipitation was performed using crosslinked protein and a GFP-Trap^®^_A Kit (Chromotek) following the manufacturer’s instructions. Proteins bound to the GFP-Trap^®^_A beads were digested using trypsin and the peptides were analysed by LC-MS/MS. Briefly, to prepare samples for LC-MS/MS, wash buffer was removed and ammonium bicarbonate (ABC) was added to beads at room temperature. We added 10 mM TCEP (Tris-(2-carboxyethyl) phosphine hydrochloride) and 40 mM 2-chloroacetamide (CAA) and incubation was performed for 5 min at 70° C. Samples were digested using 1 µg Trypsin per 100 µg protein at room temperature overnight followed by 1% TFA addition to bring the pH into the range of 3-4 before mass spectrometry.

### Quantitative RT-PCR

RNA was isolated from different parasite life stages, which include asexual stages, purified schizonts, activated and non-activated gametocytes, ookinetes and sporozoites, using an RNA purification kit (Stratagene). cDNA was prepared using an RNA-to-cDNA kit (Applied Biosystems). Primers for qRT-PCR were designed using Primer3 (Primer-BLAST, NCBI). Gene expression was quantified from 80 ng of total cDNA. qRT-PCR reactions used SYBR green fast master mix (Applied Biosystems) and were analysed using an Applied Biosystems 7500 fast machine. Experiments used *hsp70* and *arginine-tRNA synthetase* as reference genes. The primers used for qRT-PCR can be found in Table S5.

## QUANTIFICATION AND STATISTICAL ANALYSIS

### Bioinformatics analysis

Condensin complex protein sequences were retrieved from PlasmoDB (Bahl et al., 2002), EuPathDB (Aurrecoechea et al., 2010) and from NCBI databases for model organisms (File S1). An NCBI conserved domain database (CDD) search was used to identify conserved domains. PHYRE2 (Kelley et al., 2015) was used to generate 3D structure models. GROMACS 4.6.3 (Van Der Spoel et al., 2005) with CHARMM27 (Sapay and Tieleman, 2011) force field was used to perform molecular dynamics simulation in an aqueous environment using default parameters. The energy minimization was performed using steepest descent minimization till maximum force reached below 1000 KJ/mol/nm. Temperature (constant temperature) and pressure (constant pressure) equilibrium were done for 1 ns, respectively, before performing the 10 ns production simulation. Pymol (https://pymol.org/2/) was used to visualize 3D protein structure and grace software (https://pkgs.org/download/grace) was used to visualize protein stability. ClustalW was used to generate multiple sequence alignments of the retrieved sequences (Larkin et al., 2007). ClustalW alignment parameters included gap opening penalty (GOP) of 10 and gap extension penalty (GOE) of 0.1 for pairwise sequence alignments, and GOP of 10 and GOE of 0.2 for multiple sequence alignments, gap separation distance cut-off value of 4 and the Gonnet algorithm in protein weight matrix. Other parameters like residue-specific penalty and hydrophobic penalties were “on” whereas end gap separation and use of negative matrix were set to “off”. The phylogenetic tree was inferred using the neighbor-joining method, computing the evolutionary distance using the Jones Taylor Thornton (JTT) model for amino acid substitution with the Molecular Evolutionary Genetics Analysis software (MEGA 6.0) (Tamura et al., 2013). Gaps and missing data were treated using a partial deletion method with 95% site-coverage cut-off. We performed 1000 bootstrap replicates to infer the final phylogenetic tree. For orthologue identification, NCBI BLAST and OrthoMCL database search (https://orthomcl.org/orthomcl) were performed. We applied presence of conserved domain or e-value lower than 10^−5^ for protein annotation.

### ChIP-seq and global transcriptomic data analysis

FastQC (https://www.bioinformatics.babraham.ac.uk/projects/fastqc/), was used to analyze raw read quality. Any adapter sequences were removed using Trimmomatic (http://www.usadellab.org/cms/?page=trimmomatic). Bases with Phred quality scores below 25 were trimmed using Sickle (https://github.com/najoshi/sickle). The resulting reads were mapped against the *P. berghei* ANKA genome (v36) using Bowtie2 (version 2.3.4.1) for ChIP-seq and HISAT2 (version 2-2.1.0) for transcriptomic analysis using default parameters. Reads with a mapping quality score of 10 or higher for ChIP-seq and 50 or higher for transcriptomic analysis were retained using Samtools (http://samtools.sourceforge.net/), and for ChIP-seq, PCR duplicates were removed by PicardTools MarkDuplicates (Broad Institute). For the transcript analysis, raw read counts were determined for each gene in the *P. berghei* genome using BedTools (https://bedtools.readthedocs.io/en/latest/#) to intersect the aligned reads with the genome annotation. BedTools was used for the ChIP-seq to obtain the read coverage per nucleotide. For the transcriptomic analysis, read counts were normalized by dividing by the total number of millions of mapped reads for the library. Genome browser tracks were generated and viewed using the Integrative Genomic Viewer (IGV) (Broad Institute). Proposed centromeric locations were obtained from Iwanaga and colleagues (Iwanaga et al., 2012). GC content was calculated using a sliding window of 30 bp across the peak region as described previously (Lynch et al., 2010). SMC2 gametocyte sample is shown at half height due to higher level of background compared to other samples. Differential expression analysis was done in two ways: (1) the use of R package DESeq2 to call up- and down-regulated genes, and (2) manual analysis, in which raw read counts were normalized by library size, and genes above a threshold level of difference in normalized read counts between conditions were called as up- or down-regulated. Gene ontology enrichment was done using PlasmoDB (http://plasmodb.org/plasmo/) with repetitive terms removed by REVIGO (http://revigo.irb.hr/).

### Mass spectrometry analysis

Mascot (http://www.matrixscience.com/) and MaxQuant (https://www.maxquant.org/) search engines were used for mass spectrometry data analysis. PlasmoDB database was used for protein annotation. Peptide and proteins having minimum threshold of 95% were used for further proteomic analysis.

### Statistical analysis of qRT-PCR data

For selected genes identified as down-regulated in transcriptomic analysis statistical analysis was performed using Graph Pad Prism 7 software with unpaired t-test (*p<0.05 **p<0.001, ***p<0.0001 and ****p<0.00001) with standard error of the mean (±SEM) deviation.

For condensin complex subunits quantification during *Plasmodium* life cycle, statistical analysis was performed using Graph Pad Prism 7 software with Standard deviation (±SD).

### Statistical analysis of phenotypic data

Statistical analysis was performed using Graph Pad Prism 7 software using an unpaired t-test to examine significant differences between wild-type and mutant strains for phenotypic analyses; average nuclei per schizont, exflagellation centres per field, percentage ookinete conversion, oocyst number per mosquito gut, oocyst diameter, sporozoite number per mosquito gut and salivary glands sporozoites (*p<0.05, **p<0.01 and ***p<0.001). All experiments were performed in three independent biological replicates. Standard error of the mean (±SEM) was applied during phenotypic data analysis.

## DATA AND CODE AVAILABILITY

Sequence reads have been deposited in the NCBI Sequence Read Archive with accession number PRJNA542367. Mass spectrometry proteomic data has been deposited to the PRIDE repository and the original data is presented in the excel files in supplementary materials.

## Supplementary Videos

**Video S1:** Related to **Figure 6**. Ookinete motility assay for SMC4PTD parasites.

**Video S2:** Related to **Figure 6**. Ookinete motility assay for WT parasites.

## Supplementary File

**File S1:** Related to **Figure 4**. Protein sequences used for phylogenetic analysis.

## Supplementary Tables

**Table S1:** Related to **Figure 3.** Experimentally validated coordinates of the *P. berghei* centromeres.

**Table S2:** Related to **Figure 4**. List of main protein hits in the SMC2GFP and SMC4GFP co-immunoprecipitation experiments.

**Table S3:** Related to **Figure 5**. List of differentially expressed genes between SMC4PTD and WT activated gametocytes.

**Table S4:** Related to **Figure 6**. Mosquito bite back analysis of WTGFP, SMC2PTD and SMC4PTD parasites.

**Table S5:** Related to **Figure 2, Figure 5** and **Figure 6**. Primers used in this study (5’ – 3’).

