## Supplementary for "*Plasmodium* condensin core subunits (SMC2/SMC4) mediate atypical mitosis and are essential for parasite proliferation and transmission"

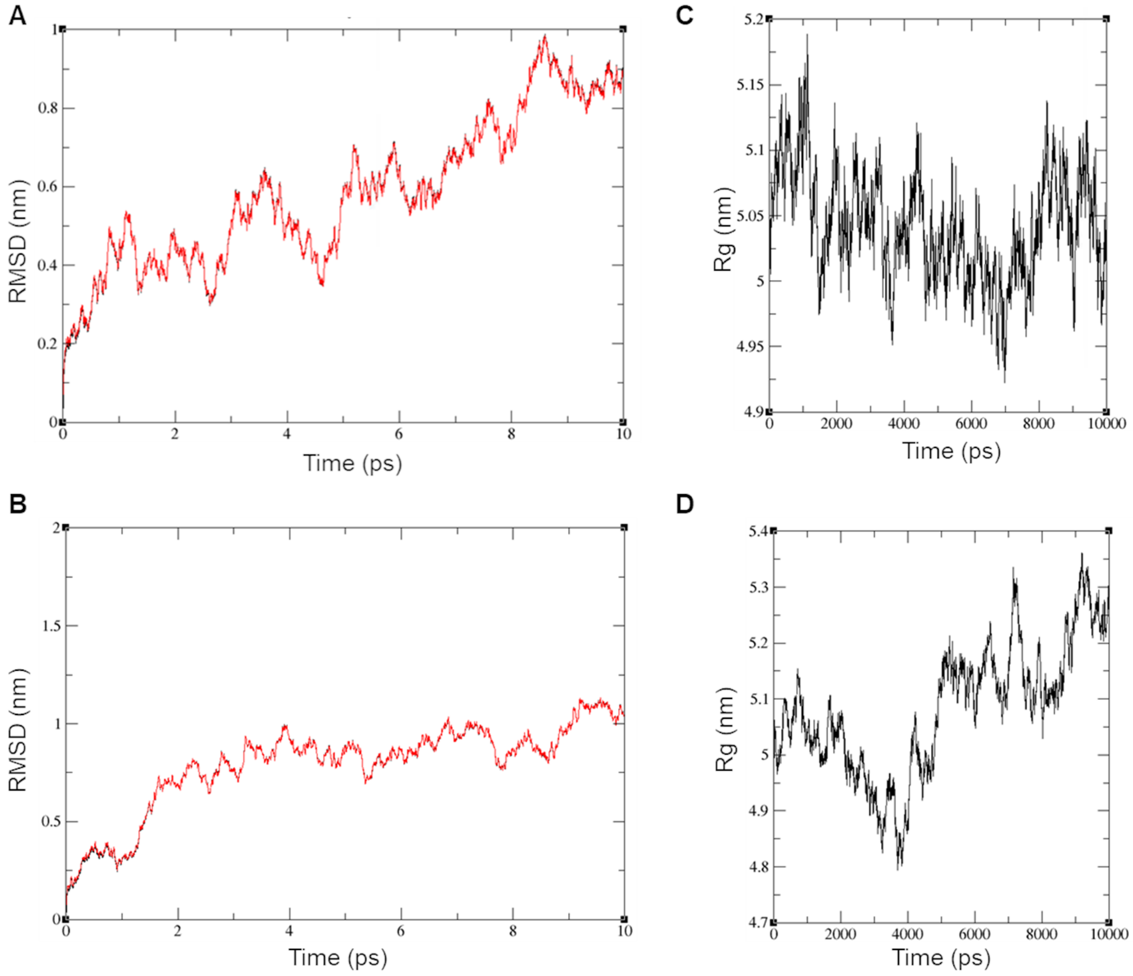


**Figure S1:** Related to Figure 1. Molecular dynamics simulation showing stable predicted 3D structure for SMC2 and SMC4. Root mean square deviation (RMSD) calculation using protein backbone structure, during the 10 ns production simulation for SMC2 (A) and SMC4 (B), respectively. Post 8 ns and 2 ns MD simulation, RMSD fluctuations becomes comparable and constant for SMC2 and SMC4 respectively. (C and D) Radius of gyration fluctuations within 2 Å suggest correct and stable protein fold for SMC2 and SMC4, respectively. Additionally, during a 10 ns molecular dynamics run the protein structure did not break, confirming stable predicted 3D structure.

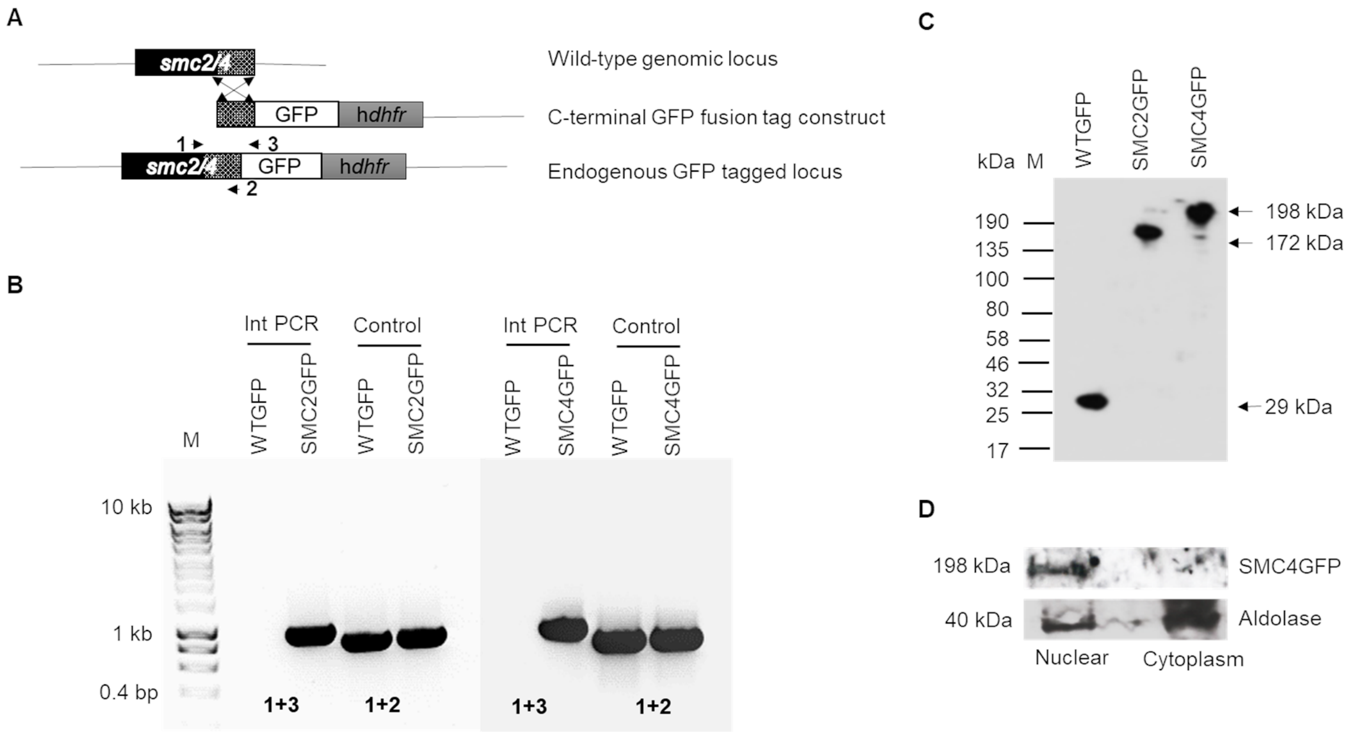


**Figure S2:** Related to Figure 2. Generation and genotype analysis of SMC2GFP and SMC4GFP parasite lines. (A) Schematic representation of the endogenous *smc(2/4)*, the GFP-tagging construct and the recombined *smc(2/4)* locus following single homologous recombination. Arrows 1, 2 and 3 indicate the position of PCR primers used to confirm successful integration of the construct. (B) Diagnostic PCR of SMC2GFP, SMC4GFP and WT parasites using primers IntT138 (SMC2, Arrow 1), IntT143 (SMC4, Arrow 1) and ol492 (Arrow 3). IntT138 and T1382 (SMC2, Arrow 2), IntT143 and T1432 (SMC4, Arrow 2) primers were used as control. Integration of the SMC tagging construct gives a band of 995 bp and 1006 bp for SMC2GFP and SMC4GFP parasite lines. (C) Western blot of SMC2GFP (172 kDa), SMC4GFP (198kDa) and WTGFP (29 kDa) protein to illustrate SMC2GFP and SMC4GFP in schizont stage extracts. (D) Western blot analysis of SMC4-GFP for subcellular localization from schizont stage extracts using anti-GFP (nuclear) and anti-aldolase (nuclear and cytosolic).

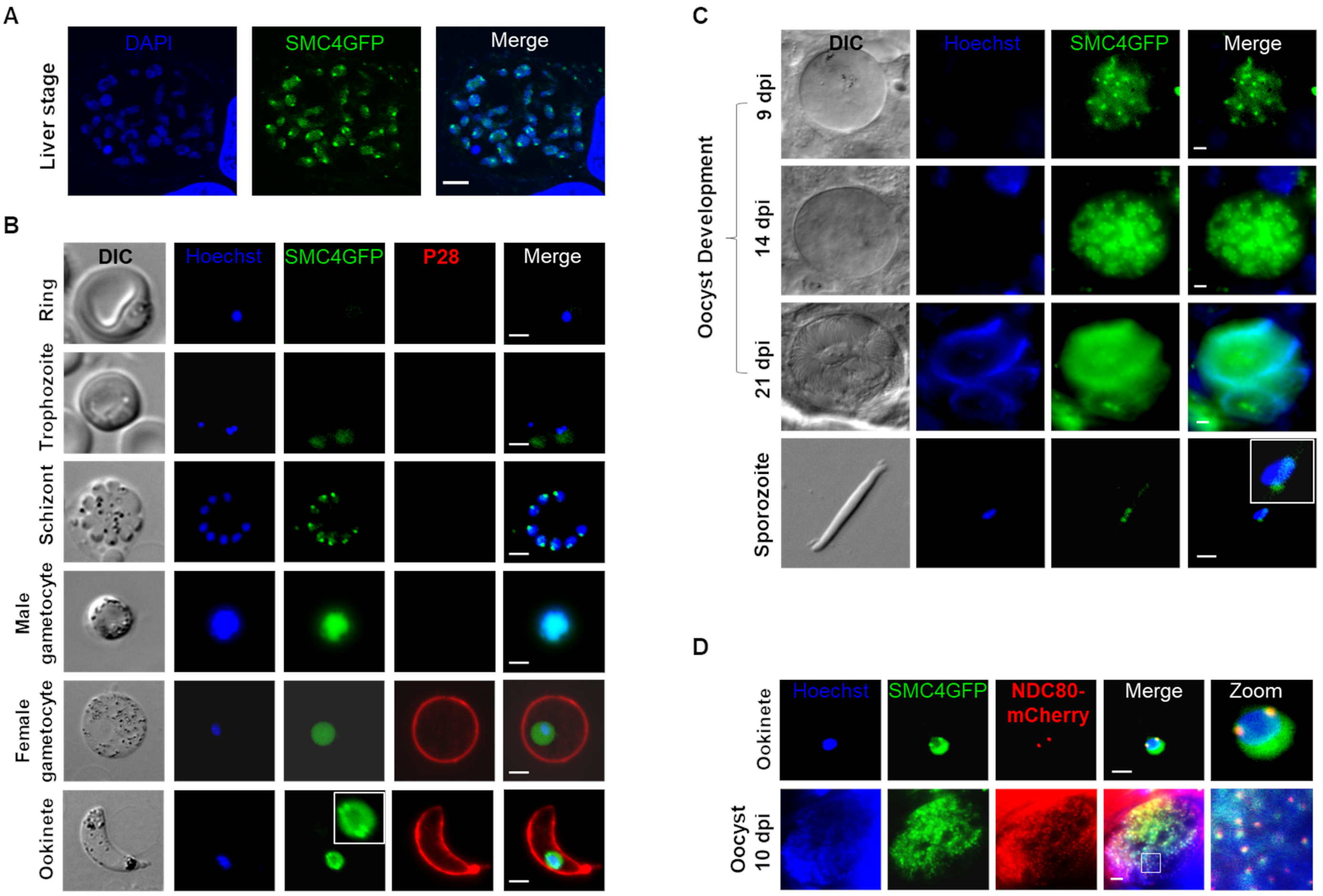


**Figure S3:** Related to Figure 2. Localisation of SMC4GFP throughout the *Plasmodium* life cycle as detected by live cell imaging. (A) Liver stage schizont at 60 hours post infection. Merge: DAPI (blue) and GFP (green). Scale bar = 2 μM. (B) Asexual blood stages and sexual stages at different time points. Merge: Hoechst (blue, DNA), GFP (green) and P28 (red, cell surface marker during female gamete activation, zygote and ookinete stages). 100X magnification. Scale bar = 2 μM. (C) Sporogony in the mosquito oocyst; 9 days post infection (dpi), 14 dpi, 21 dpi (Scale bar = 5 μM) and mature single sporozoite at 21 dpi (Scale bar = 2 μm). 63X magnification. Merge: Hoechst and GFP. (D) Live cell imaging of SMC4GFP and NDC80mCherry localization in ookinete (Scale bar = 2 μm) and oocyst 10 dpi (Scale bar = 5 μm). The white box represents magnified region. Merge: Hoechst (blue), GFP (green) and mCherry (red). 63X magnification.

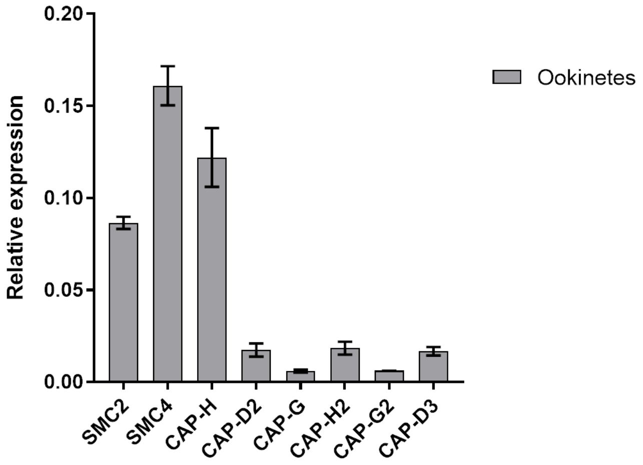


**Figure S4**. Related to Figure 4. qRT-PCR analysis of condensin complex subunit expression in ookinete stage of parasite life cycle. Error bar = ±SD, n= 3. Primers list has been provided in Supplementary Table S5.

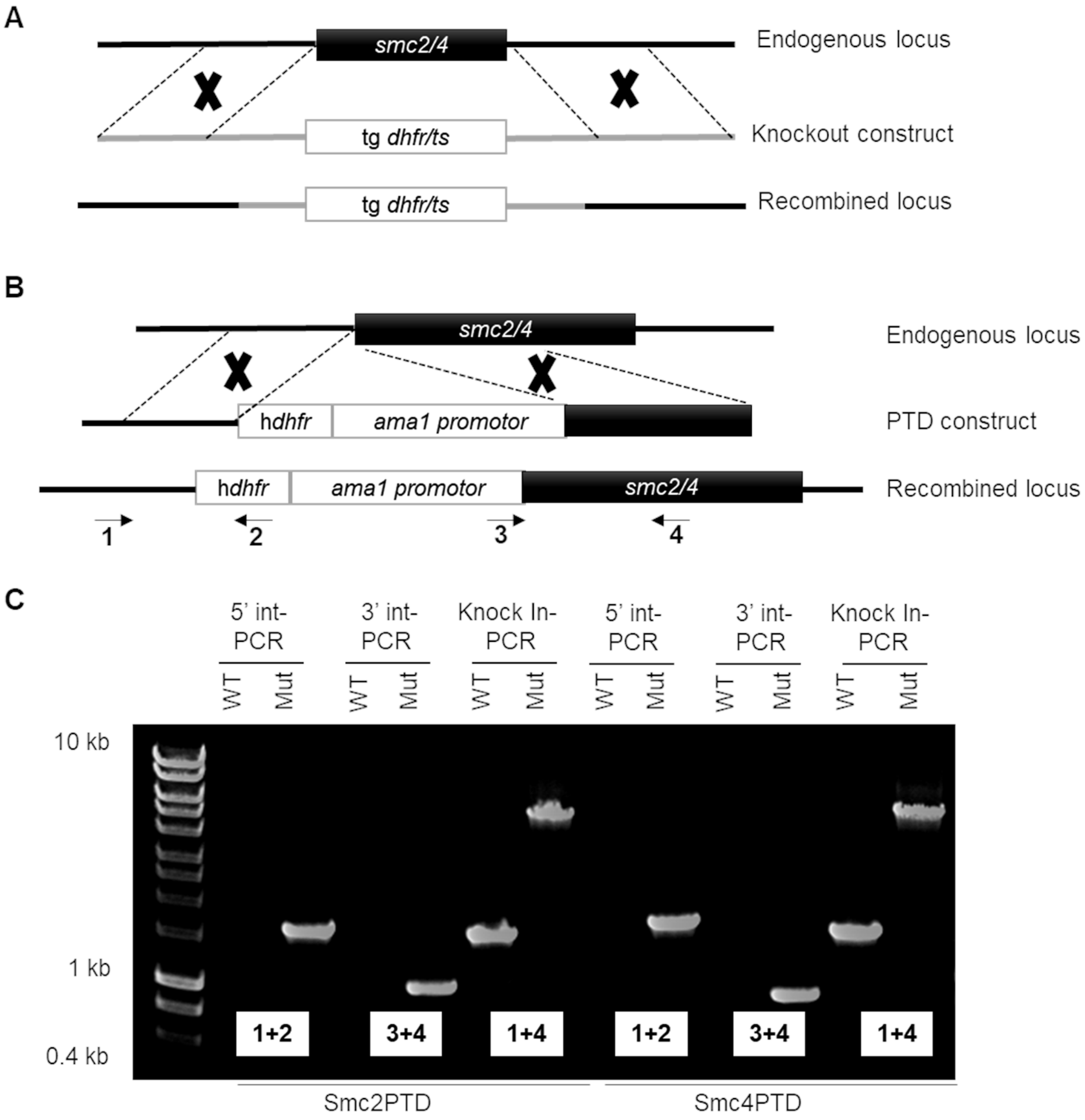


**Figure S5.** Related to Figures 5 and 6. Generation of knockout and conditional knockdown of SMC2 and SMC4 genes using a *dhfr* drug-selectable marker or *ama1* promoter trap double homologous recombination (PTD), and genotype analysis. (A) Schematic representation of the endogenous *smc(2/4)* locus, the targeting gene deletion construct and the recombined *smc(2/4)* locus following double homologous recombination. (B) Schematic representation of the promoter trap strategy (SMC2PTD and SMC4PTD), placing *smc(2/4)* under the control of the blood stage *ama1* promoter by double homologous recombination. Arrows 1 and 2 indicate the primers position used to confirm 5' integration and arrows 3 and 4 indicate the primers used for 3' integration. Primers 1 and 4 were also used for Knock-In PCR. (C) Integration PCR of the promotor trap construct into the *smc(2/4)* locus. Primer 1 (5'-IntPTD18 [SMC2], 5'-IntPTD008 [SMC4]) with primer 2 (5'-IntPTD) were used to determine successful 5' integration of the selectable marker resulting in a band of 1518 and 1655 bp for SMC2PTD and SMC4PTD, respectively. Primer 3 (3'-intPTama1) and primer 4 (3'-IntPTD18 (SMC2) and 3'-IntPTD008 (SMC4)) were used to determine the successful 3' integration of *ama1* promoter resulting in a band of 1004 bp and 877 bp for SMC2PTD and SMC4PTD, respectively. Primer 1 (5'-IntPTD18 and 5'-IntPTD008) and primer 4 (3'-IntPTD18 and 3'-IntPTD008) were used to show complete knock-in of the construct with a band at 4698 bp (SMC2PTD) and 4653 bp (SMC4PTD), and the absence of a band at 1538 bp (for SMC2, endogenous) and 1493 bp (for SMC4, endogenous) resulting in complete knock-in of the construct.

**File S1**: Related to Figure 4. Protein sequences used for phylogenetic analysis. Protein sequences can be retrieved from EupathDB (https://eupathdb.org/eupathdb/) and UniProt (https://www.uniprot.org/) database.

>sp|G5EGE9|DPY26_CAEEL Condensin complex subunit dpy-26 (Kleisin Iγ) OS=Caenorhabditis elegans OX=6239 GN=dpy-26 PE=1 SV=1

MDVPSSSNVTGRRKRQVLDDDEDDGFRSTPLRKVRGTKKIRPADVVPETIMTKIGAHIDDIVNKKKVG
ELNCFEYKSPLEIHTIEDMIKAKASIQEMAVVLEGAQCIIGYRVDRLHHDVRQIDSALSSGTVMRDSNG
EEIHLTLESRKAKKKMAVVDGMNGMLDFLNNMDDALTTTELDADNDKNWKEDEENIAGEPRIDFKAN
SKDVDAFLQRDIFPEKLIYALSIKRATDLRADLLSDVSNYISADDTAHDLKDANIDWLRANPTFQKATKG
SVCNSSNSFHSLNYYGIHSPDGRTLMLHNRIADKNADDRFFTSDVSVSLVKNTRALLTNSLDKKPRIL
DNYLMLEVKDRPVIGRYKIMSKDVKKSTLPLAESSREKDLANLTFAEMNHRPSNLDMTVAGASDMSM
LPGNQGLPLAQGENDETIALDRLTPPLQSSVSQKASDEYVLPPLEANDLDEHLIGKLPIEPNEMDQTL
ANMFDKKLEVFNTSDTLESKVWKNGIRAEEWGEDDEAIMKNDTKHPRQAGIEGWIKATDAWTNYDV
VKMNVNREARSQLDENAIDEQESYRNMVPEIGKNLFLVKSDDYMNNYPGDRPADFTVNDEVSDVMK
MWSGEDSTAEDDVPLEQIQQEIREQVQQQDVDMEPIEEMDYDAGGAAFDVDFDDRLAAPVEVEEM
EGDNNRNDGRVADILFNEQMDETEVEERNEQDVQRELEDIALAADEVAELMTSAPPPQLVGPSAEM
REEIQNIGKNDNAHWVPPVVGDQERQAAVTAQRKRREKKAKSRKATVEDFVHYFRDIPDDEIEREITA
AKCSKIADEKSTFLSEQQLYLPTLGIENKPHVAFEMGLLGNSGMFFKKSYGKIRLERVKNQKAEQDLFI
DEARGNKDSDCLNWLLSFSGFRCMENPEPITGSDSDENLRTAVEQPFDDDFANDYYDEDRYDPNYE
QQLAAAQMGPDMQRKLALTASHINQMFPNIHSKRYGGEYGDSDDEFDDSFDRQSIQAKNLDAAKHK
KCLAEILKTDSLSMPSIQYVLEQLTSNQTLRMNNTTIRAADDRNETGRPATPTMEADKTLTSVFDYRS
PNKSNHDVNETMKALTEMPDYQAADERPNNQPTTSTYGTANTENRKVHINGCHTLLSLALSMPSRM
GETVRPSSIVSFLLHIANENNLQIVQDRSKRSWMSDFIVLNSSESLPRGLKMGRIEDQDEFWKRTQDP
DAIEGTASDANNVFSNLMRRPKAVPVRKGRGAGGQPTTSDLGAIVEEEEMEE

>sp|Q15003|CND2_HUMAN Condensin complex subunit 2 ((Kleisin Iγ)) OS=Homo sapiens OX=9606 GN=NCAPH PE=1 SV=3

MGPPGPALPATMNNSSSETRGHPHSASSPSERVFPMPLPRKAPLNIPGTPVLEDFPQNDDEKERLQ
RRRSRVFDLQFSTDSPRLLASPSSRSIDISATIPKFTNTQITEHYSTCIKLSTENKITTKNAFGLHLIDFM
SEILKQKDTEPTNFKVAAGTLDASTKIYAVRVDAVHADVYRVLGGLGKDAPSLEEVEGHVADGSATE
MGTTKKAVKPKKKHLHRTIEQNINNLNVSEADRKCEIDPMFQKTAASFDECSTAGVFLSTLHCQDYRS
ELLFPSDVQTLSTGEPLELPELGCVEMTDLKAPLQQCAEDRQICPSLAGFQFTQWDSETHNESVSAL
VDKFKKNDQVFDINAEVDESDCGDFPDGSLGDDFDANDEPDHTAVGDHEEFRSWKEPCQVQSCQE
EMISLGDGDIRTMCPLLSMKPGEYSYFSPRTMSMWAGPDHWRFRPRRKQDAPSQSENKKKSTKKD
FEIDFEDDIDFDVYFRKTKAATILTKSTLENQNWRATTLPTDFNYNVDTLVQLHLKPGTRLLKMAQGHR
VETEHYEEIEDYDYNNPNDTSNFCPGLQAADSDDEDLDDLFVGPVGNSDLSPYPCHPPKTAQQNGD
TPEAQGLDITTYGESNLVAEPQKVNKIEIHYAKTAKKMDMKKLKQSMWSLLTALSGKEADAEANHRE
AGKEAALAEVADEKMLSGLTKDLQRSLPPVMAQNLSIPLAFACLLHLANEKNLKLEGTEDLSDVLVRQ
GD

>sp|Q9Y7R3|CND2_SCHPO Condensin complex subunit 2 ((Kleisin Iγ)) OS=Schizosaccharomyces pombe (strain 972 / ATCC 24843) OX=284812 GN=cnd2 PE=1 SV=1

MKRASLGGHAPVSLPSLNDDALEKKRAKENSRKQRELRRSSALHSITPRRESLNNSSPFNSSHQVPV
LSNFEEWIKLATDNKINSTNTWNFALIDYFHDMSLLRDGEDINFQKASCTLDGCVKIYTSRIDSVATET
GKLLSGLANDSKVLQQTEEGEDAENDDEDLQKKKERKRAQRSVKTLVKDFESIRAKKFELECSFDPL
FKKMCADFDEDGAKGLLMNHLCVDQHGRIVFDSSDTVIKDLENKDVEAESQEAVVAAPIESHDTEMT
NVHDNISRETLNGIYKCYFTDIDQLTICPSLQGFEFDSKGNLDVSLLKSLSDEVNMITTTSLVDNTMEKT
DADAASLSSDSDGEEGHIVHALEEMAYDEENPYVDVVPKAMDESENPDFGVDTEVNMADGSTMNE
NYSIISTAAANGVYEYFDKSMKKNWAGPEHWRIQALRKNINNASTVFNSSNTAESSDNVSRSLSSTE
RKKRRELDNAIDFLQEVDVEALFTPATSSLKLPKSHWKRHNRCLLPDDYQYDSKRLLQLFLKPKMSVL
PNADGEGQLQLNKALDDENDLDGIQPHGFDSDGSDNVDEGIPPYGFGDSDSPKQTPLLTPPSSSGF
GDNLLLTARLAKPDMLNYAKRAKKVDVRVLKEKLWKCLDLENTIKENSINSHIEGSEMESEETNMPVK
SFFSTVNQLEETYEKKELKDISTSFAFICVLHLANEHNLELTSNEDFSDVFIRPGPNLTTLEALENDV

>sp|P38170|CND2_YEAST Condensin complex subunit 2 (Kleisin Iγ) OS=Saccharomyces cerevisiae (strain ATCC 204508 / S288c) OX=559292 GN=BRN1 PE=1 SV=3

MTTQLRYENNDDDERVEYNLFTNRSTMMANFEEWIKMATDNKINSRNSWNFALIDYFYDLDVLKDGE
NNINFQKASATLDGCIKIYSSRVDSVTTETGKLLSGLAQRKTNGASNGDDSNGGNGEGLGGDSDEAN
IEIDPLTGMPISNDPDVNNTRRRVYNRVLETTLVEFETIKMKELDQELIIDPLFKKALVDFDEGGAKSLLL
NTLNIDNTARVIFDASIKDTQNVGQGKLQRKEEELIERDSLVDDENEPSQSLISTRNDSTVNDSVISAP
SMEDEILSLGMDFIKFDQIAVCEISGSIEQLRNVVEDINQAKDFIENVNNRFDNFLTEEELQAAVPDNAE
DDSDGFDMGMQQELCYPDENHDNTSHDEQDDDNVNSTTGSIFEKDLMAYFDENLNRNWRGREHW
KVRNFKKANLVNKESDLLEETRTTIGDTTDKNTTDDKSMDTKKKHKQKKVLEIDFFKTDDSFEDKVFA
SKGRTKIDMPIKNRKNDTHYLLPDDFHFSTDRITRLFIKPGQKMSLFSHRKHTRGDVSSGLFEKSTVS
ANHSNNDIPTIADEHFWADNYERKEQEEKEKEQSKEVGDVVGGALDNPFEDDMDGVDFNQAFEGT
DDNEEASVKLDLQDDEDHKFPIRENKVTYSRVSKKVDVRRLKKNVWRSINNLIQEHDSRKNREQSSN
DSETHTEDESTKELKFSDIIQGISKMYSDDTLKDISTSFCFICLLHLANEHGLQITHTENYNDLIVNYEDL
ATTQAAS

>sp|Q8C156|CND2_MOUSE Condensin complex subunit 2 (Kleisin Iγ) OS=Mus musculus OX=10090 GN=Ncaph PE=1 SV=1

MRIPRSETMNSSFLKARGQQDVLSSPLERVPPASRPGKAPLGTPKTPVLEDFPQNDDEKERMQRRR
SRVFDLQFSTDSIHLASPNRNIDVSTTISKFTNTQITEHYSTCIKLSSENKITTKNAFGLHLIDFMSEILKQ
KDAEPTNFKVAAGTLDASTKIYAVRVDAVHADVYRVLGGLGKDTPPQGEESHSGDGSTLETERTKKP
AKPKKKQSCKTIEQNLSNINVSEADGKCAVDPMFQKTAASFDECSTTGVFLSTLHCQDYRSELLFPSD
MQTLSSGEPLELPDLGFVDMTDLEASLQQCVEDRPLCPSLAGFQFTKWDSETHNESVSALVDKFKK
NDQVFDINAEAEDDEEDVPDGPLVEDFVDNDEPDLSAAGDHEEFRSWKELCQVQSNQEEVISLEDR
DIQVMCSFLSMKPGEYSYFSPRTMKMWAGPDHWRFRPRPKQDATSCTEHKKKSAKKDFEINFDDDI
DFDAYFQKTKAATILTKSTLENQNWKATTLPTDFHYETDNLIQLHLKPGKRSLKMDQDQKAKTEHYEE
IEDYDYNNPNDTSNYCPGLQAADSDYEEADDLFADPVGTLDLESDPKTTQENGHISPENQGVDITTY
QELNLVAEPQKVNKIEIHYAKTAKKMDMKKLKQSMWSLLTKFSRKEADTEANHTESGQEGAPEEVAD
EKKLSGLTKDLQTRLPPLMAQNLSIPLAFACLLHLANEKNLKLEGTEDLSDVLVMQGD

>tr|P91663|P91663_DROME Condensin complex subunit 2 (Kleisin Iγ) OS=Drosophila melanogaster OX=7227 GN=barr PE=2 SV=1

MTLPRLETPLRRSAVGSYQEGVSRMLTPFNDDEAERREARRRTLLQQHHRSSTLESIEDNETIKNCL
ELYNGNKVSKDNAWNLMLIDSLANLLDHHHKRMSNFKMAGSSLEASSKVYGLRVDSIYLDAMRISAG
LSARTLTDKQINAAEDDDGPQGEQATGEGQDSAQQAAKEAAPKPKRQKKPISTVTKNRETLNSRLDT
APLQDPVFGKLNSTCGASINASNRLMHNILPSFDSELRLRTTYNFWNSEESTEEVQDHTTLNAEMEQ
WPATSLMSTNLMRKLLPHAERSNLRPLHTGYIITSAPNPKSANEKAAEVVQDEDHDEGLDNADDVCV
NKISMAFDINAECEPMPDLDGPPPLVLEVDSNELEELTAEEQMVINNCRRLRKQTEFIEDLRPVDGNS
KLEYSYRPMDQISQFWAGPSHWKFKRTRPRSTFSQTNGQVDTQPIRTQRAKKSAHLNANRRAKALD
YGNVTENFFQQLDTTIRQRKANFQKKWDPRKLILPTKFELDPDLFFKYESAPSIKLSKRAGEPDSDEG
GDLGIDMDADMHHDDDNDQELFNNEHFTDAVPANVSVIAAIAAEQAAEASMMNVSAGEIGLTQMNAT
CNNTVFEIGTEFEGAPSQVAKVIVPFAKRAKVIDMKNLKKSCNSLIQKQLLNAVPEETIPSHPKKKGEH
YSKGFASFQQVYQKLPDLLTTKMSDSLSPSVAFYAVLHLANDLKLRLIPQEDLEDFQIRQVLD

>sp|Q564K3|CND2_ARATH Condensin complex subunit 2 (Kleisin Iγ) OS=Arabidopsis thaliana OX=3702 GN=CAPH PE=1 SV=1

MDESLTPNPKQKPASTTTRIQAPTSPFFLGSNDDRLEREQARAARAAASRRRSVIFARGSQPETESD
PCFDKQQILELFQNCIKLASENKINQKNTWELNLIDHLCEIIKVEDENNTETNFQKASCTLEAGVKIYSM
RVDSVHSEAYKVLGGITRAGHDDGGDHEDAAGAVENATNQKKQPEKKISPLSTLEPSFDALNVKKFD
VAFAVDPLYHQTSAQFDEGGAKGLLLNNLGVYGGCQVLFDSQEIPGKLVSSANKHDKSETIDLSFVK
ECVEQMVLNMRKKDEIVPSLRAIINQFDEENQRPSDTFSCGQQTTESFDISHGNDASYADDDEGYEN
FGTSFDYEGQSGDVDENFGPNEAEPIYSNFHEEVEPASLQDMDSDDRLENVDDYLFLSLGISSKQNS
WAGPDHWKYRKTKGPDVQPASEIKSSPPAKKTRKKKQAEPELDFAKALEEEMPDIFAPPKNPKTLLL
PASRTPCQTKLPEDCHYQPENLIKLFLLPNVMCLGRRRRKNSGETSRQQPDDYEHGESWGNDNVY
DDDDGPFDDNENDQSDAEDTNTLISQPRQVNKIDVQYDKASKQVDVQVLKETLWECLQESHQPPIQ
DEEHQQEPPESRSFKVLLASFPDDCQAAERTQDISPHLCFICLLHLANEHNLSLIGSQNLDDLTIHLA

>sp|Q8BSP2|CNDH2_MOUSE Condensin-2 complex subunit H2 (Kleisin IIβ) OS=Mus musculus OX=10090 GN=Ncaph2 PE=1 SV=1

MEDVEVRFAHLLQPIRDLTKNWEVDVAAQLGEYLEELDQICISFDEGKTTMNFIEAALLIQGSACVYSK
KVEYLYSLVYQALDFISGKRRAKQLSLVQEDGSKKTVNSETPCETENEFLSLDDFPDSRANVDLKND
QASSELLIIPLLPMALVAPDEVEKNSSPLYSCQGDILASRKDFRMNTCMPNPRGCFMLDPVGMCPVE
PVVPVEPYPMSRSQKDPEDAEEQPMEVSRNGSPVPVPDISQEPDGPALSGGEEDAEDGAEPLEVAL
EPAEPRTSQQSAILPRRYMLRERQGAPEPASRLQETPDPWQSLDPFDSLESKVFQKGKPYSVPPGV
EEAPGQKRKRKGATKLQDFHKWYLDAYAEHPDGRRARRKGPTFADMEVLYWKHVKEQLETLQKLR
RRKINERWLPGAKQDLWPTEEDRLEESLEDLGVADDFLEPEEYVEEPAGVMPEEAADLDAEAMPES
LRYEELVRRNVELFIATSQKFIQETELSQRIRDWEDTIQPLLQEQEQHVPFDIHIYGDQLASRFPQLNE
WCPFSELVAGQPAFEVCRSMLASLQLANDYTVEITQQPGLEAAVDTMSLRLLTHQRAHTRFQTYAAP
SMAQP

>sp|Q6IBW4|CNDH2_HUMAN Condensin-2 complex subunit H2 (Kleisin IIβ) OS=Homo sapiens OX=9606 GN=NCAPH2 PE=1 SV=1

MEDVEARFAHLLQPIRDLTKNWEVDVAAQLGEYLEELDQICISFDEGKTTMNFIEAALLIQGSACVYSK
KVEYLYSLVYQALDFISGKRRAKQLSSVQEDRANGVASSGVPQEAENEFLSLDDFPDSRTNVDLKND
QTPSEVLIIPLLPMALVAPDEMEKNNNPLYSRQGEVLASRKDFRMNTCVPHPRGAFMLEPEGMSPME
PAGVSPMPGTQKDTGRTEEQPMEVSVCRSPVPALGFSQEPGPSPEGPMPLGGGEDEDAEEAVELP
EASAPKAALEPKESRSPQQSAALPRRYMLREREGAPEPASCVKETPDPWQSLDPFDSLESKPFKKG
RPYSVPPCVEEALGQKRKRKGAAKLQDFHQWYLAAYADHADSRRLRRKGPSFADMEVLYWTHVKE
QLETLRKLQRREVAEQWLRPAEEDHLEDSLEDLGAADDFLEPEEYMEPEGADPREAADLDAVPMSL
SYEELVRRNVELFIATSQKFVQETELSQRIRDWEDTVQPLLQEQEQHVPFDIHTYGDQLVSRFPQLNE
WCPFAELVAGQPAFEVCRSMLASLQLANDYTVEITQQPGLEMAVDTMSLRLLTHQRAHKRFQTYAA
PSMAQP

>sp|Q9LUR0|CNDH2_ARATH Condensin-2 complex subunit H2 (Kleisin IIβ) OS=Arabidopsis thaliana OX=3702 GN=CAPH2 PE=2 SV=1

MTSHGGGEVRGERIHTVQPERDLVANWEVDLSEKLEEYLLKICSGEITGNEEDGQIPVNFAEAALLLQ
GSVQVYSKKVEYLYNLVLRTLEFLSKQRDQEQSKGTSNENEASSSRQVDEEENDLFWNVDDIPVDT
KNRLDSSVGGDTCPSQFVKPPANLVVLEGDCLDTSGDGGELESYLLATTHLYRDFILLDPCDAVAVN
EFLGDNYGGKGRNSAHRGSSVRKSFHSSVGRSGGSARKSSVGKNQGTNVHLSPICGNGPNDQNC
DQGSQPPVFEDNDHGFDMDNEYGGAMDFSDTDADEDDPWKPLNPYEPGKLKVKPFKKVKILKKIG
WSITKDHMTSMFPLARPNGPISSELIEIWKMHGCASKDEQASQDIPYYEKLREMLVNGGNQPCGANG
NYNDNDKDNHDEANNGDFHDFGEHDGDDAEHPFMDEDVLNMNDGGAAEFHNYDGFENGESNCQ
ESLEDLCRSHLDALLANIAKSEKQTDLAARVSTWKQKIEQNLEEQELHPPFDIQEYGDRIINKLTVEES
GNVETFTDLMKDQEKHEVARAFSALLQLVNNGDVDLEKPGNSTNEPMCYTAVKPFSVRLLKVHNRK
NEKRGIHLPQKRAKSPITKGKSHESPPPKKRNTCSVSSQTRKVSLKISKINGVGVRCTPNSKKRRKGR
SDDVTEVTEVASIEKSLGKL

>sp|P34341|KLE2_CAEEL Condensin-2 complex subunit kle-2 (Kleisin IIβ) OS=Caenorhabditis elegans OX=6239 GN=kle-2 PE=1 SV=2

MTRNAPPGQESTDLAWLVTPAKDLVENFSIDVLKALAGYLEVIRQESEDTDNQVDAATTYRLFDFQR
ACRIIQGSCAVYGRKVDHVYELTISVVDLVENKGQDDGNTGSRRGAGRRKNFNLGSTNYDLADIDSL
KQEALANFEKTVKEEKKSIDAVRMVENAEVIESQYERKSCLVAKPTQFMFKLNYGQLNRTDEQILNAK
SRPDVIGKVKDFEIKKSKVKHDQQILYSHDCYRGNLDQFTLPGARWMPDNKELAANFGVADLEVELD
LEQEHEKISAYGPFKDPLSGREVVPPPRWFIEQEAVRQNQEIQSRATSRAITIAAKTLRDSQGFGSQP
TRLSQPFVERHRQSNHLNDFLSFVEGRVNKNRPSTHLTTGLVDMFVDNFGSVMQNDEPNTSRRPDE
NYAPMDFDDDFGGGGDDDDDDYIRNLSRRDEKRAPAPWDELDKNHIIWYTGDENLPVVSKPVKKITK
FQPKPAEMLARKQRREEKINKSRRDEFMETHDYLQDYYYWRSAARINPIKDWKIESLRTAILAEKKRR
IKEKTAKIREARIQNMQRKRTARVIPVEQFEPVTEDIPTSNRRTLGAEYDDVVDEDLAAEVELSMFGG
GFDDDEEDVRPRGERPPMAPNNLEFDALQTDFDIPPAEYVPLRFEDIDDAELNSVINLPGNLLIDKALP
LLKKFAENRTDREQMAYEMAKAYEDVDVAVSTLQEHVDKWHSRMEPILEEGETRKEYDVHAVGRAV
IGQYDIEGGTKRLLDLVMDRPWYEISRYFLSCLFMCNVGNVMVSEDMELPLEERINSMKITLLKRDMH
CEMFKEAGALDA

>tr|Q8INL2|Q8INL2_DROME Chromosome associated protein H2, isoform E (Kleisin IIβ) OS=Drosophila melanogaster OX=7227 GN=Cap-H2 PE=4 SV=2

MERILPEEAEAAYLSEAREQILEIAKNRPGTQVAKCIRAYDEQQDLASLVVLLEELSKNTDYSTDLRISL
GYYIEELLRHCLGRNDVSRRSAIVAAGSALQYCGKIYGDRVEYLCQVVEHQIEALLTSELQKETPSGS
AAPEKNERRPEETRKRHPKKLTNKEVDPYLLTLEPKRFKTMSDDKRFNAAGFVKCTRNRTIEYLYQD
HTPPNLWKHAPIVDPHNPYDQDEKKQYKMFTYHVEHRYNTLLPDIPFERLNLIKEYVHTNQVNTTEIL
NEHMTTKEYLDEYIALENQMLAARYGAIVTRRRRLVDSARFMEDNLDEGLAKKMCMDKNLPMDTNE
TVLIDQSLVDENSRSLTTAESTMGISQAENSTLKSSEVEATLSVSHIENSLSDSQEENPPLSSTLAINES
SVLDSTRVDPIKDLTLDELLLIDSGISMEELSDIQMHTAGQSFDDEGVVLSDLEDQRQLSPMLQMVSP
SMEAKTLNIIEMDADLIMNVPREVSYPILLNVMGLPIKRLRRKCIFKLPPEFDLFRQARLPIKREGQQKS
PTTPRTVLQIGREAPQNEREPGSPCSLEFDEDLNFLGFRQRRPTFDSGFDIEEPRVSSCVSTIGEVKT
ELEDEIEANTNNLTDASASDALNENVNDTKESGLGDSLAQELNASTEPNAENNTANSSQLEVTATESS
GTLSGLETTVDLGMESALVSSVLDLSTEPSAMDSNIINVTQPDNSDLEISTVQDQDTLHDEADANDLS
AETPNPIADSAVDDCSLIRDWHRRLAPALEAAHERQNFNIKDLGTEILDICKAGNRTATLADVMADKDP
SIMCRYMLASLVLTNHGNVSLDFENRDKSKPIDMSQFRMHLKSMKRMEINPEDDVGNINAAQSKSTP
RSKQLNDSAHKPRTSTTASAPTKRKSAENSLSEVFAKTVRLIQPIPKMWPTPSDADSGISSMGSSLAS
TARLK

>PF3D7_1304000  | Plasmodium falciparum 3D7 | condensin complex subunit 2, putative (Kleisin Iγ) | protein  | length=1024

MKKLGVNNAGENKGNSIQNKDGTNKKTIEVNKNMRRLTFLNNESNDSIEGNKNKYDKNKVKEINDVF
KNCMVALSHNKICTRNAFDIHIIEHLEDLINLNDEEIPEELNDEMIENGEFNLSFTRASKAIEGATKVYGY
RVEAIYDQTYNFLTNMNLAKQFELDNDMMDDNKNTIDPLNKRMRKRKLTYLQESSTLAKSSDITVDSL
SLSNISVDTFFLKLNSTYDHSCGQKYLLPNLNLNNDLSIQFDGDIDVCEYKKRRKTLDDIGKDGVKDKE
DGIHIIMKNKNDDNCDTYDKIICNNNDINKVDEKKNNDVDICENRNKKNESCNIYVDTFNVNEHKEFFDI
YKKKLFLNADILKEILYGGPDDFNSLHICPELDYFKEELKKHKLKRTDSKDTDDVDKEDGDGNLYNDD
DIGNTDKNNIINFYDENLGCNMSNNMLMESIYNSDALNTDRKNIFPENLGMSCGNLNEEKKNNNMNS
SNNNLNNCNYDNNDLYLGDYRMDDLNIENVMQESLGFDNLNYSCDKNMNFSNNNGMLLQQSINMM
DGMNFPELIKSENKELDLGNTQINNEHYLLKNNYDIHKRHSIISEVPDDDTLWNRSVMTFENRINAIDIN
NELNYHYYMPNKLMMMGNFRNLIDINKNVTNNINNNNKSTILQNIIMQKKMKYAFDVTSIDFENLYTEN
NNIELSTYDLWKKEKKKYVSNALFCIDQTSYIFDTRENCVNCVNTVTDKVMKFSKFCVYPDFLHACNK
NRNKHNNNNINKNHMNVILNEVNNDNILNDIDPINQDDFNNNIDNFDMENMDDMNDGHLHEGLNEAI
DKYYNMDFDNIWANNDNKNNDNINNNNNLNNSTYNNMNSHNNNNNNNNNNLSLFKKHSNFFNTALF
QFNHSNTFGNVPFENVSKFVDVAKIKKILCDIVKPNEEQKSNTNDSLCEYQTEKDNQIVTYTAEKTTTF
EEIVKETTAKLNESEASSTSIHMLFVCLLYTCNDQELLLEKIPNQNNFYVRYGLPVECHVNHDDIPMLK
N

>PBANKA_1402500  | Plasmodium berghei ANKA | condensin complex subunit 2, putative (Kleisin Iγ) | protein  | length=943

MKKLGVSNKTNTNFTQIKGPDPIFKNPIEFNKNLRRLSFLNNKDEDLNKSEKNKVKEINDVFKNCMAAL
SHNKICTRNAFDIRIIDHLEDLVNLNDEEINEELNDELLETGDFNLSFTRASKAIEGATKVYGYRVEAIYD
QTYNFISNMNIAKKSETNDDVIDEKKHANEITNKKVKKRKLEFFQESSTLAKPSDITIESVSVSNISVDTF
FLKLNITYDHSSGISYLLPNLTLNNDLSIQFDGDIDTCEYKKKMKFEEKLGKQNIERRNEKNCGSNDEL
NDMLQKNESPVKVGEYVTFNDNDKIMAREYKSKLYTNSDILREMFFGNEIEEFNNLNICPELDYFKEEI
KNLKLKRSDSKILDDEDIDNYNDDDTGLDKFSKKKNELNLDGNMSMDNLLGDMDGNNNDDGNNEHM
NDNANYGDNNKILESSFNNNLNFDDCNIDDLNIENVMQESMVFDNMNLNDSLNNNLNLSQNILSLHH
NSNILGSSIPLPELMKSENKDFSLINSFTGNFNFSSQNMLFKNQDKGSPSKPKNMLSIIPDDDTLWNR
QAISFENRLNAIDVNSKFNYHYYNPSKLMINGNFANLMSMAKVAFKNKQGPLNALTNKKLKTSFDITYI
NFENLYIEVNDVELSAYDLWNKEKKKYISNSLFAIDQTSYIFETKDNCINCVNTVIDRIMKFARSPFIESQ
NFNTDIKTNVILNEINNDYIGNDLDQINNFERKQSENMFMDNMDDGQDYQMHEGLNDSIDKFYNMDF
EDIWQNENKNDITKFGSKNDNTSIFQLHQSMGHTNSLGSVIAPDNLPKFVDVSKIKKILFNIVKPDENE
ENVENGKSEENGKSEENGKSEENEENKKNSDSSKQIVPYEGEKTTTFKSIINKTKTKLTESEVNGTSI
HMLFVCLLYTCNDQELLLEKIENEEDFYVHYGLPVEFHIKQDDTRMLKN

>PY01534  | Plasmodium yoelii yoelii 17XNL | CCAAT-box DNA binding protein subunit B (Kleisin Iγ) | protein  | length=886

MKKLGVSNKANANFTQIKGPDPIFKNPIEFNKNLRRLSFLNNKDEDLNKSEKNKVKEINDVFKNCMAA
LSHNKICTRNAFDIRIIDHLEDLVNLNDEEINEELNDELLETGDFNLSFTRASKAIEGATKVYGYRVEAIY
DQTYNFISNMNIAKKSDTNDDVIDEKKHVNEITNKKMKKRKLEFFQESSTLAKPSDITIESVSVSNISVD
TFFLKLNITYDHSAGISYLLPNLTLNNDLSIQFDGDIDTCEYKKKMKFEEKLGKQNIERRNEKNLGSNDE

LNDMLQKNESPVKVGEYVTFNDNDKIMAREYKSKLYTNSDILREMFCGNEIEEFNNLNICPELDYFKE
EIKNLKLKRSDSKTLDDEDIDNYNEDDDTGLDKFSKKKNELNLDENMSMDNLLGDMDGNNNDDCNN
ENMNDNGNYGDNNKILESSFNNNLNFDDCNIDDLNIENVMQESMAFDNMNLNDSLNNNLNLSQNML
SLHHNSNILGGNIPLPELMKSENKDFSLINSFTGNFNFSSQNMLFKNQDKGSPSKPKHMLSIIPDDDTL
WNRQAISFESRLNAIDVNSKFNYHYYNPSKLMINGNFSNLMSMAKAAFKNKQGPLNALANKKLKTSF
DITDINFENLYIEVNDVELSVYDLWNKEKKKYISNSLFAIDQTSYIFETKDNCINCVNTVIDRIMKFARSP
FIESQNFNTDIKTNVILNEINNDYIGNDLDQINNFERRQTENMFMDNMDDGQDYQMHEGLNDSIDKFY
NMNFEDIWQNENKNDITKFGSKNDNTSIFQLHQSIGHANSLGSVVAPDNLPKFVDVSKIKKILFNIVKP
DENEENAENENEENKKNSDSSKQIVPYEGEKTTTFKNIINKDANFVNRAHELYICILYIYFLT

>PVX_122040 | Plasmodium vivax Sal-1 | hypothetical protein, conserved (Kleisin Iγ) | protein | length=928

MKKLGVNTKAGANLTRIKENENLFKKPIEFNKNLRRLSFLNNKNNDANQKDHQDKCDKTKVKEINDVF
KNCMVALSHNKICTRNAFDIRIIDHLEDLVNLNDEEINEELNDEMLETGDFNLSFTRASKAIEGATKVYG
YRVEAIYDQTYNFLSNMNIAKQSEVNEELVEEKKNANEISNRKIKKRKLEFLQESSTLAKSSDITMDSV
TVSNISVDTFFLKLNSTYDHSSSNSYLLPNLILNNDLSIQFDGDIDACEYKKRKKMEEATEEGVDEDTA
ATQTSGKFPPMIRSSSSYDNDCSDAMVKCYKKKLFLHSDVLRDILFVAGNEDFNSLNICPELDYFKAEI
SKHKMKRSESKELEDGEQGNDEDDDDDDEEDDRRYAYNLNEEEHMSMHKGDDYGGERAKNNLAS
SVGGNLTMDNLLDDCDFHNASVGDGRMLNSSVNNNSMHFNDYKIEDLNIENVMQESLAFDNMNLND
SVGNNLNYSQSILSFQQGSNTMTGLPLPELMKSENKDFSLMNSLGGNFPLSTQNSLFFKNQGGSPT
RKGLISTIPDEDTLWNRNVMTFENRLNAIDVNNKFNYHYYVPSKLMAHGNFRNLMDIAKGTHKNKHM
VLQSVVQKKVKASFDVSLIDFENLYKEINDVELSTYDLWKKEKKKYVSNALFSIDQTSYIFETKDNCINC
VNTVTDRIMKFSKAPMIAPGDYCGDLKLNVVLNEVNSDFVTNDMGQIHYGEGRNMESTFPENMDEH
QDCHMQEGLNDAIDKFYNMDFDDIWQNENEHNLSKQGSKNENASALQLRQRVSHGTTLGNANMEN
VAKFVDVSKIKKILCDIVKPAGKEATGSEGPAQRNDQIVPYVEEKTTTFKDIIIATKSKMNPEEVNGTSIH
MLFVCLLYTCNDQELLLEKIPNEDNFYVRYGLPVEYHVKPDDMLMLEN

>TGME49_288930 | Toxoplasma gondii ME49 | hypothetical protein (Kleisin Iγ) | protein | length=1185

MVARATDGGDTGAPAGSETHPVVRLGFRPDGKRRLSVVSGPASLSTGDGAPSGESRRGVGGTTGA
SLAGHAPGALARQASSLNSASSSFSSFASSSAFPASLPFSSSANRPGFARPTARAASGAGRALAGRL
LVRAFFDDMKTVSMQRINQKNAFQVDLIDRLALVVHQQLVKTDNAVLTDGTDGQAPALLDGASSPDL
SSTSELSRTGRGEEGEEGITFTHVSSAVEGATRVYGYRVEAVYDQTYHVLNLMSSSRQGQGGEDAG
DGDAAAHPTRRGRHHQQLALFKKGGASTLAPASEITESQIEKDSCVDPYFLKISGMFDQAGAKGLLLA
NLEVDTSLRMKLDGECRAFPTGVLRNGREKGNAAREETEANHAPEETAEVAEPSVDCAFLRDLLLAG
ETPASVLALDICEKPIGHFRELLQSLRRGRDVKAGDALSEAEEETNELDVDDDMLDVDMQGTTLGDS
GFQQATQEDEELLKIGESYMDDEAASGGLREVLEAQEAESREVTQDDELFGDGVDFFCDGHVGDAD
DDRDGSWDRPRGEIRERDAEGERILYEVAADESSASSCTHPKSYSFDRLVPAFSQMLGVPAGGHGA
QLAAKRAFGLVPTAAGSQGGGDPEGRETGRRRGSGEETSSSGCVDFFPLSAKGSAAAVGRPSAPS
ARLTVKERQLRKQEQFREMLDPFRVDMSNLDTKSGGLPLGSKFQCYVPKPQENVTVASDFTSSPFL
HSPHLLVSLAMVPSKEIRLLRHTGPSVGDGWAPDGAGNSGGRAGRDLGRGDSQAPDGPWEATFID
AGDDGGLEDDSFWSAAAHGPLLKESALEAADAKAQTLLEGGGGLDRDWSDTAWAGGENDQASSYI
AHLFERGDAASRGGAAGLSLGDIVLLAEPQKVGVSELRLSIPSRYVDVAVVKKALKTALGVVPDFSAD
QDAENSALRRKRKRDMLTAPGEEEPQEEDDVDSCLDSVDGETKEPCRDLALWNPRGRRDGIDFETL
CAETTRKLPSTEKANLSPQMLFVCLLYTCNEETLHLEQSADFSSFSAFVHAPTDWHMRDDAEAVRVL
DTVRHYTPLALPAVPEKKKKETLSLEAPSAESAADDGGAAGDADDAQSPAKRRGALGDEERRQKRR
REGRGGKHEEGNRKGHRDEETGSKKRRGKKRRKEDSDESSDGSDDEEAAASEDRDGSC

>cgd3_3960 | Cryptosporidium parvum Iowa II | Condensin complex subunit 2 (Kleisin Iγ) | protein | length=901

MARLSLHPRIFESEYSAETDNYLEENVSNLNKNGWSGRKIQKSVLPNKRRSISIGQSRNDDSETTRLS
RTSINLKGQIDQNKSFFSKSSSDQLPNEELDKLCNQCLNLLRQNKISSKNAFDILLIDHLNDIVNVQDSQ
NEEEKSNIIKKENKENVSKINNNSKEKNIEIDKSKMNKGEFHNEMISSETTSQDTNFQKFQRAAVTLEA
SARIYGYRVDSTFDNAYRILSNIKSGQILARNDSDCEEDEKDQQEDDSSGNRLEESSKKKKYKRNLIF
SGENNTIVSNPESITLKEFEKTKEMSEKPFFVNFLNDKVKIDNIRFGLDYNNIGSISSMLMNNLELKDSL
YNRDKSASLTFNISGRNNLINSNNFSCSTICKDELFKVNKKTLDLLLPNPNENFGKDISNIVCPNLSRIF
DSIQEITENYGDIKQVKDSTSIEEYQLNEQVIRRFFEFTDPSFENDSLESENNFLKKHSQSMDDIEYVN
ENQDLEFLNNFEEEEKVELNMGLQDFQMHVERVIKDLNKTESSEHILEDLDQNKDIEIEKKSGEFKFS

KIGDVLDFLAKKSTKSLEFFNERKKGEKSNSVFGQKLRSKSENTNFEIPSPLLTNNIFNGIEWMNNLPK
IKQTQINSSRSKTKNLSSNKLFVGKNSTTSIFNYNLNHLFCLANLTGVKINLRSIHKAENSEETAGQNYI
TNDILIKTNNSQHINKNILNNSMSEQTDLMVNHCEIEDLMQVESYLQPLSQEEFSIDNGKDILIQSKLLLP
FDEDKISDTDILLEKSSKSLHVRDEYNSSLKFSKSSNHVDIDLIKNVLKNSIQILQSDDSNSLTLFNIINQS
RKLLQQDGLNSVSTGIFFICILHICNENNYSLNLSDQNANSDSPIELNSENYVHIMLNLNSSVDLQK

>ETH_00039510 | Eimeria tenella strain Houghton | Ccaat-box DNA binding protein subunit B, related, related (Kleisin Iγ) | protein | length=122

GFDEEEPLEAAAAAAAERVSFCHLSSAIEGATKVYGYRVEAVYDQTYHLLNGLSSLKNSSNGVEDDE
EQQQQQQQGSKRKQRQQQQLALFFKGGSSTLADPQDILNEQQDTGVLIDPFFV

>BBOV_I002180 | Babesia bovis T2Bo | hypothetical protein (Kleisin Iγ) | protein | length=726

MASRKRSRDVDSSEISEAPQPVSKKVTGPSGNTQDLLTLFTDCMSALSTNKICSRNAFEVGIIDHMTD
LVHLDDGSVDDDVVELLPEDTSSGASRRLNFTRASKVVESASKIYGYRIEAIYDQTFNVLMSMNSANQ
ADGTSGSTSATKPRGRHRVKIDLTTSSRTLAPESEVTLTEIPMDNVILDPYFLKISSMFDHSGAMGLLLI
NLQVTDDLSLDLDGDSLVFPPPRARSEHDDSNVILSRDAVKKCFFGNTEPKRMEILPEAAYFRQELER
LREMRRRKECGDVIASDSDGEINPEPESKHIDFFKVRQPIAPIEDSLEPMDAMGHIDDVPEDTVHDMPI
SGTPPDTGSDTGSVPNSTGIALRQRLAEIDLIGGSQFSYYTSVPVKTHVPSSKNKGSDTEPDDTNLKT
QRPVKSRQNITNDLESYLRNIDLEEKIASVELSTVFTTPKAVGKKAPSTFAASDFTGAIYRFNDTFLTRL
GLLGNRCLRFVSDHEWRNGPNTDSSVPILQVHYCADIDAPYSTMRISENFSWRLDEQVGLLDEDDM
WNASNDFEDPELPLTQDGPVVANIESMYESEPGEPLLALSQHADPAAISQWPREGAIVPAPVAAYVDI
FKIKKTLCGVIVPPPAFHEALEDKQQIDKEHGSGNTQSEYSCGFQDAVTDTVSRLMDSDVAALTSHIL
FVCLLHVCNEQDLLLKQSRPLEDFVICAGAPKEQHLGDVQGSL

>PBANKA_030180| Plasmodium berghei ANKA | condensin-2 complex subunit H2, putative (Kleisin IIβ) | protein | length=775

MSTQEDEVRLLIQNLQKCNNTNECINFDLASTIQEFLNSLDKNSFEDIDKTIRENEKDKDLMNSFTSAAI
FLENCVKILGLKIEHLHNLAHNTLYNIYKENKNSNSNKKQLLIIDEEEYLYINEIKNLKNTITENDIIEEDLL
VKTIPLPTFLFTDHIRVKNKTENHKNIIKYDEKDNFLINKKKLLNEPDIENIPYIDTLGNESINSIETKIMDLN
STNSYKNMDNLNLIDNKQTIEISSVNSLNFDKLFLENDGIILLDINDYNIFINDEYDFTLQNKNSTILFEKY
EFFSRNSIYLSNNLTEYIHEQNTIQHTYKINNIYDITSLRLCTDTLLFKTDFYSYDIALDIIKNKNYLINKFE
RQKKKLYILDETIHKDHKYNTIYKQNADYCDYCGSIITSIIEPNEPNNRCIYCNNNTQIDRNEKNGLYKKL
PGYYNLSCYNITETEDFLTYMQPNKIIDIIMKNEINDINLDSNSTTNKFEENIIDPIFHQNNDFTDKGDNY
RLSNDQKLFRQIKIPSLYIQKLGLNIDYYYLEPLIYNLIKNLKKEKNVDRFFSINFYDHNENYDIEILKDDY
YHEIKDEQNKTIQETLNMDTFINIKSIDNHVKNFPTSILKKTDSNTSLVFSFEDKIQDRVNAWSNFLEEK
LVILKRQPQYNVEYYKKKILKYIINNGDNIYFPDLIKNDEKYQIYRNFLTTLMLINTNKLNISEIDQQKHSN
NITNYQINIKNINVNEYMNISSAFDNTKFTINDKKRKITEKSDNIDNSFHLEKKNHI

>PFB0185w | Plasmodium falciparum 3D7 | condensin-2 complex subunit H2, putative (Kleisin IIβ) | protein | length=797

MSTTDELNLLIQNLQKCNNTNECINFDLSSTIQGFLNCLDRNVLENIDKGLGENEYEKEVVDNFTSAAI
FVENCVKIFSQKIEHLHNLAHNTLYNIYKENKHNSSSKKNQLIMSDEEEYLYINEIKNMKNTQHDNDIIE
DDILIKTIPFPTFLFSDNIKKTKDINEDKRKTNFNNNEEDKEKDNKNKDNDIDAINEFEITDNNSVNTLNF
EKIFIENDGILLLDINDYNVFIDDPYNFSIQNKNSTILFEKYDFFSRRSTYLSSNTLSKYVVENKNMDHIY
KLYNHITDIINKNICFDIFLFKQDFFDYDFSLGILKNKKSILNKFKQQQKKLHPLEENTHMDTHHINNNHH
LQKYDLNRPLPNYYMLHCYNIKNYQDFFRYMQPNYILEIMKRHIIKEIYNTNQQERAIQKEAYEIYNEQT
KKKNDHKENNNIDVPKYKDNTKCYDSPFYNYYISNNIIQFDHLIDDDMIYFDEYFYKSLILYNTNINDLHK
NTNNNQTNDETNIINNMKDEKQKNLIIYSNINNFSNDQKLFNQIKIPELYIQKLGLNFSYYHLEPLIYNFIK
TLKKKNDFEKFFSVNLFDDKPIYEFDILRDDEYDEQKNEDNKNHIEENINFENITDKNILNDEMDNIPIAI
FENDHLDNTFIMNDDQELQDRVSKWNAFLEEKLEILKRQPKYDLDLYKKNIINYTINNGENILFTKLIKN
KDKFEISRNFLTTLMLINADILNIKKINKHKKSNNISNYEIHIKKENLQQYLSISKQVQNKSFLIKEKKRKK
NKQHLTNGMKDTSKKKQKI

>PVX_003630| Plasmodium vivax Sal-1 | hypothetical protein (Kleisin IIβ) | protein | length=784

MSTPDELTALIQNLQKCNNTSESINFDLASTIQEFLNCLDRNVLEELEGGAHEGEKERESDREREKEG
EREREQDVANSFTSAAIFVENCVKIFGLKIEHLHNLAHNTLYNIYRENKHSNAGKKHMVMSDEEEYLFI

NEVKNLKSCPGESEPCMEEDLLVKTIPLPTFLFSENVKKKEGAAEVDSELGIDLSADLDTPRRSDNGS
GTDDRRGHSNLSKESEEAAREANGETDSIASSPAWRKGENDRLTEEAEEVLEQNSLKPLNFDKMYL
ENDGILLLDINDYNVFINDEYDMSILNQNSSMLFEKYDFSCSRHSTYLSPANLTKYIEREKTVGDIYRTY
HFNDILSDDLCSDVFLFKSDFSHYDLALGIIKSKKYLLSRFKEQKKYLYVLDENAHMDKDRVGTSTIEK
NDIKRKLPHYYMLNCQNIKRAQDFFTYMQPSQIIDIITRNARRRYLSCLPSGETSQGGQSGNAAGADS
PVGGAASHTSPPATPAPPPPGRSTDQKLFEQIKIPDIYVQKLGLNFRYYHLEPLIYNLIKELKKKKNVEK
YFSLNLFDEQNNYDFDILQDEEYADEQNEGNAAAEGEGNLTDENFLDVRSLNGDMVGGMMGGMMG
GMMDDIPLDVFEKKDSSDAFFASFDEDIHDRVNKWNAFLEEKLQLLRSHPKYDVDQYKKNIIHHTLNS
GAKTPLCNLIKDREPYQVCRNFLTTLMLINTNMLQISEVNQHSQSNDVSNYQINVKKENVQEYLGSSK
RFKNASFAIKDKKRKTASKGGTRNAKPAKKKPHKD

>PY01684 | Plasmodium yoelii yoelii 17XNL | hypothetical protein (Kleisin IIβ) | protein | length=773

MSTQEDEVRLLIQNLQKCNNTNECINFDLASTIQEFLNSLDKNSFEDIDKTIRENEKDKDLMNSFTSAAI
FLENCVKILGLKIEHLHNLAXNTXYNIYKENKNNNSNKKQLLIIDEEYLYINEIKNLKNTITENDIIEEDLLIK
TIPLPTFLFTDHIKVKNKIDNHKNIIKNDEKDNSLLNKKKNLNKSDIENIPYIDTLENESITSIQKNVMDLNS
EYSYQNMDNMDNMNSMENKQTLEISSVNSLNFDKLFLENDGIILLDINDYNIFINDEYDFTLQNKNSTIL
FEKYEFFSRNSIYLSNNLSEYIYEQNTIQHTYKINNIYDITSLRLCTDTLLFKTDFYSYDLALDVIKNKNYLI
NKFERQKNKLYILDETTHKNHKYNTIYKQNANYCDYCGSIIEPNESNNHCIYCNNNNEKNGFYKKLPG
YYNLSCYNITETQDFLTYMQPNKIIDIIMQNEINDTNLDANSTTNKLDQNIIDPIFHENNDSSDKSDDQKL
PNDPKLTNDQKLFKQIKIPPLYIQKLGLNIDYYYLEPLLYNLIKSLKKEKNVDRFFSINFYDHNENYDTEIL
KDDDYQEIKDEQNKTIQETLTMGTFINIKSIDNHVKNLPTSILKKTDSNTSLAFSFEDKIQDRVNKWRNF
LEKKLDILKRQPPYNVEYYKKKILKYMISNGDNIYFPDLVNDNEKYKIYRNFLTTLMLINTNKLDITEIEQN
NSNNITNYKINVKNMNVNEYINFPNSFDNIKFTINDKKRKITQNFDNKNDSLYLQKKHHI

| Experiment | | Bite back (dpi) |
|---|---|---|
| 1 | WTGFP | 4 |
| | SMC2PTD cl 5 | - |
| | SMC4PTD cl 3 | - |
| 2 | WTGFP | 4 |
| | SMC2PTD cl 5 | - |
| | SMC4PTD cl 3 | - |
| 3 | WTGFP | 4 |
| | SMC2PTD cl 5 | - |
| | SMC4PTD cl 3 | - |

**Table S4:** Related to Figure 6. Mosquito bite back analysis of WTGFP, SMC2PTD and SMC4PTD parasites. dpi = Days post infection after mosquito bite to mice.
